# A burst in T cell receptor translation mediated by eIF3 interactions with T cell receptor mRNAs

**DOI:** 10.1101/2019.12.20.885558

**Authors:** Dasmanthie De Silva, Lucas Ferguson, Benjamin E. Smith, Grant H. Chin, Ryan A. Apathy, Theodore L. Roth, Marek Kudla, Alexander Marson, Nicholas T. Ingolia, Jamie H. D. Cate

## Abstract

Activation of T cells requires a global surge in cellular protein synthesis, accompanied by a large increase in translation initiation^1–4^. A central component of the translation initiation machinery–the multi-subunit eukaryotic initiation factor 3 (eIF3)–is rapidly turned on when quiescent T cells are stimulated^3^. However, the precise role eIF3 plays in activated T cells is not known. Using a global transcriptome crosslinking approach, we show human eIF3 interacts with a distinct set of mRNAs in activated Jurkat cells. A subset of these mRNAs, including those encoding the T cell receptor (TCR) subunits TCRA and TCRB, crosslink to eIF3 across the entire length of the mRNA. The *TCRA* and *TCRB* mRNAs do not co-localize with translationally repressed environments of P-bodies or stress granules but form distinct granules, potentially acting as translation “hot-spots.” T cell activation through CD28 causes a burst of TCR translation controlled by elements in the 3’-untranslated regions (3’-UTRs) of the *TCRA* and *TCRB* mRNAs that directly contact eIF3 and that are required for T cell activity. These results highlight a new role for eIF3 in regulating the translation dynamics of the TCR and provide insights that can guide the engineering of T cells used in cell immunotherapy applications.

## Main

Translation initiation serves as a key gatekeeper of protein synthesis in eukaryotes, and requires the action of eukaryotic initiation factor 3 (eIF3). In humans, eIF3 is a 13-subunit protein complex that coordinates the cellular machinery in positioning ribosomes at the start codon of mRNAs. Several lines of evidence indicate eIF3 also serves specialized roles in cellular translation, by recognizing specific RNA structures in the 5’-untranslated regions (5’-UTRs) of target mRNAs^1^, binding the 7-methyl-guanosine (m^7^G) cap^2^ or through interactions with *N*-6-methyl-adenosine (m^6^A) post-transcriptional modifications in mRNAs^3^. Binding to these *cis*-regulatory elements in mRNA can lead to translation activation or repression, depending on the RNA sequence and structural context^1–4^. These functions for eIF3 can aid cell proliferation^1^, or allow cells to rapidly adapt to stress such as heat shock^3^. Additionally, eIF3 plays an important role in the development of specific tissues^5–7^.

In the immune system, T cells are activated by contact with a foreign antigen in a process that involves a burst in protein synthesis^8–11^ that correlates with assembly of eIF3 subunit EIF3J with the main eIF3 complex^10^. However, whether eIF3 serves a general or more specific role in T cell activation is not known. To delineate how eIF3 contributes to T cell activation, we first identified mRNAs that directly interact with eIF3 in Jurkat cells activated for 5 hours with ionomycin and phorbol 12-myristate 13-acetate (I+PMA), or in non-activated Jurkat cells as a control, using photoactivatable ribonucleoside-enhanced crosslinking and immunoprecipitation (PAR-CLIP)^1,12,13^ (**Fig. 1a**). In the Jurkat PAR-CLIP experiments, RNA crosslinked to eight of the thirteen eIF3 subunits, as identified by mass spectrometry: subunits EIF3A, EIF3B, EIF3D and EIF3G as seen in HEK293T cells^1^, as well as subunits EIF3C, EIF3E, EIF3F, and EIF3L (**Extended Data Fig. 1a-c, Supplementary Table 1**). Consistent with its role in T cell activation, eIF3 crosslinked to a substantially larger number of mRNAs (∼75x more) in activated Jurkat cells compared to the control non-activated cells (**Extended Data Fig. 1d-f, Supplementary Tables 2 and 3**). Notably, in activated Jurkat cells eIF3 interacted with a different set of mRNAs compared to those originally identified in HEK293T cells^1^ (**Extended Data Fig. 2a and 2b)**. These mRNAs are enriched for those encoding proteins central to immune cell function (**Supplementary Tables 4, 5 and 6**). Interestingly, the extent of eIF3 crosslinking in activated Jurkat cells does not correlate with mRNA abundance (**Extended Data Fig. 2c**), suggesting that eIF3 is involved in specific regulation of T cell activation.

**Figure 1.**
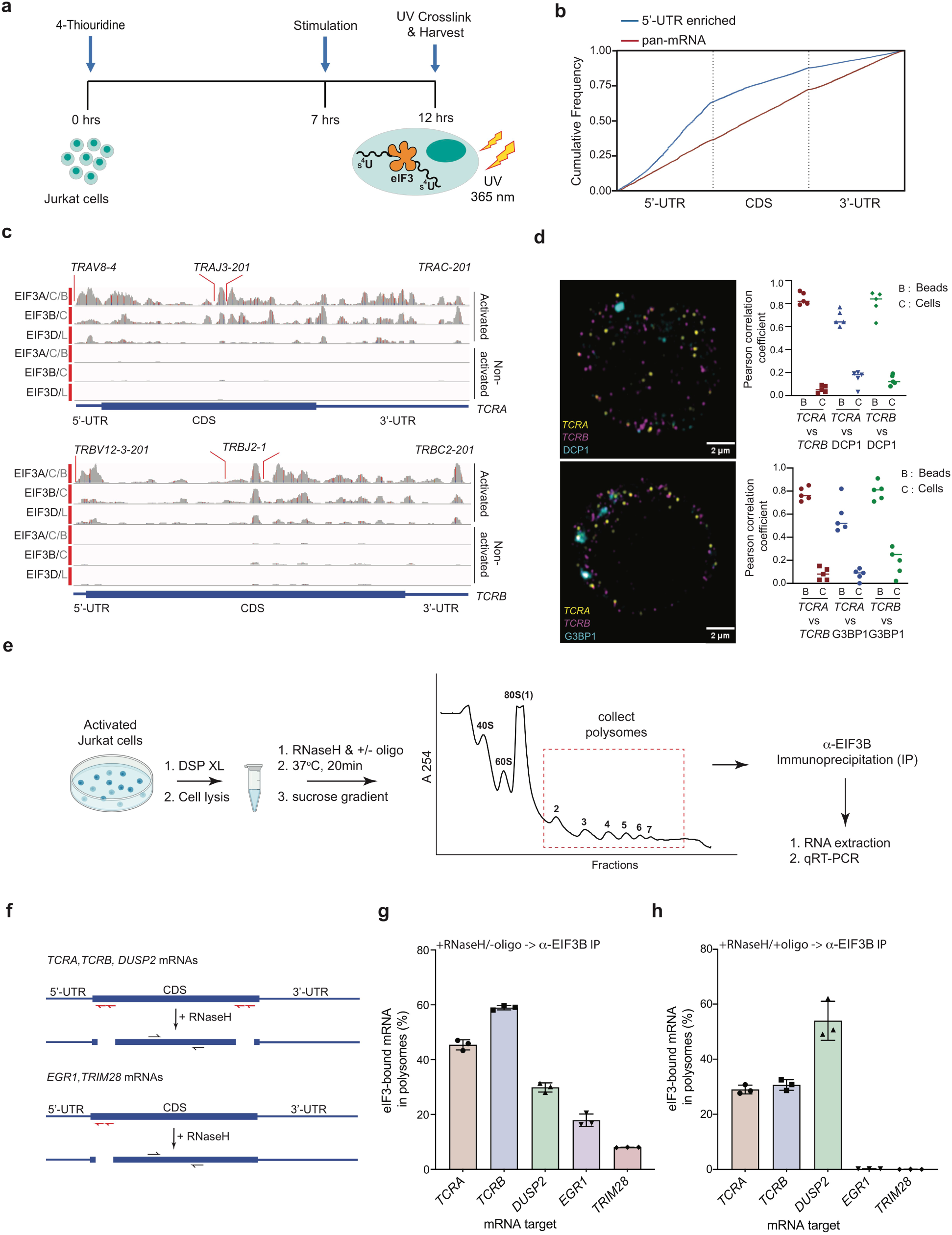
eIF3 interactions with mRNAs in activated Jurkat cells. **a**, Schematic of Jurkat cell treatment with 4-thiouridine, activation, UV crosslinking and cell harvesting for PAR-CLIP analysis. **b**, Varied mRNA crosslinking patterns to eIF3 in activated Jurkat cells. Cumulative plot showing mRNA crosslinking in sample EIF3A/C/B to predominantly the 5’-UTR (n = 396, 414 mRNAs in replicates 1 and 2, respectively), and across the entire length of some mRNAs (“pan-mRNAs”, n = 634, 621 mRNAs). The 5’-UTR, CDS, and 3’-UTR regions are shown normalized by length. **c**, Crosslinking of the indicated eIF3 subunits EIF3A/C/B, EIF3B/C and EIF3D/L across the entire *TCRA* and *TCRB* mRNAs in activated and non activated Jurkat cells. 5’-UTR, coding sequence (CDS), and 3’-UTR elements (below) along with the variable (V), joining (J) and constant (C) regions (above) for the mapped TCR genes in Jurkat cells are shown. The blue and red vertical lines in the plotted reads indicate the amount of T-C transitions vs other mutations, respectively for a particular nucleotide. The *TCRA* and *TCRB* mRNAs are present in both non-activated and activated Jurkat cells (**Supplementary Table 4**). **d**, FISH analysis of *TCRA* and *TCRB* mRNAs (yellow and magenta, respectively) and P bodies (on the top) marked by the location of DCP1 (cyan) and stress granules (on the bottom) marked by the location of G3BP1 (cyan), in activated Jurkat cells. Graphs to the right of the images indicate Pearson’s correlation coefficients (PCCs) of *TCRA* and *TCRB* mRNAs with each other or with P bodies or stress granules. Labels on the x axis are, B: TetraSpeck™ microsphere beads, C: activated Jurkat cells. (*n* = 5, P <0.008, for PCC values of cells relative to bead colocalization, across all the channels tested, using the Wilcoxon rank-sum test). Images are representative of one experiment of the five independent experiments in the graphs. **e**, Schematic outlining RNaseH-based assay of eIF3 interactions with mRNAs in polysomes. DSP refers to the dithiobis(succinimidyl propionate) crosslinking agent. Oligos, DNA oligos designed for RNaseH-mediated targeting and cleavage of specific mRNAs. See the Methods for details. **f**, Strategy for detecting mRNA fragments released by RNaseH digestion. Red arrows denote DNA oligos for RNaseH-mediated targeting of mRNAs. RT-qPCR primers (black) were used to detect the CDS regions of the mRNAs. **g-h**, Amount of eIF3-bound mRNA co-immunoprecipitated by anti-EIF3B antibody, from polysome fractions of lysate treated with RNaseH either in the absence of mRNA-targeting DNA oligos in panel **g**, or in the presence of mRNA-specific DNA oligos in panel **h** (red arrows diagrammed in panel **f**). The percentage is relative to the amount of total mRNA present in the polysome fraction prior to the IP. All of the IP experiments in panels **g** and **h** were carried out in biological duplicate with one technical triplicate shown (*n* = 3, with mean and standard deviations shown).

In activated Jurkat cells, eIF3 crosslinked to mRNAs by a multitude of patterns (**Fig 1b-c, Extended Data Fig. 2d-f**), consistent with varied roles for eIF3 in T cell activation and function. Many of the mRNAs have a single PAR-CLIP site in the 5’-UTR as observed in HEK293T cells^1^ (**Fig. 1b, Extended Data Fig. 1f, Supplementary Table 3**). However, eIF3 crosslinked to some mRNAs across the entire length of the transcript, from the beginning of the 5’-UTR through the 3’-UTR (**Fig. 1c, Extended Data Fig. 2e**). This “pan-mRNA” pattern of eIF3 crosslinking–which includes polyadenylated mRNAs as well as histone mRNAs–has not been observed before. Interestingly, a number of these mRNAs encode proteins important for T cell activation, including both the alpha and beta subunits of the T cell receptor (TCR), subunits TCRA and TCRB (**Fig. 1c, Supplementary Table 3**).

Crosslinking in PAR-CLIP experiments requires direct interaction between the RNA and protein of interest^14^. Thus, the pan-mRNA pattern of crosslinking between eIF3 and certain mRNAs suggests formation of ribonucleoprotein complexes (RNPs) highly enriched in eIF3. Notably, the pan-mRNA crosslinking pattern in the *TCRA* and *TCRB* mRNAs occurs in activated but not in non-activated Jurkat cells (**Fig. 1c**), suggesting eIF3 may contribute to translation activation of these mRNAs. Since Jurkat cells have a defined TCR, we were able to design fluorescence in situ hybridization (FISH) probes across the entire *TCRA* and *TCRB* transcripts. Interestingly, both the *TCRA* and *TCRB* mRNAs formed distinct puncta in activated Jurkat cells, but did not co-localize with either P bodies or stress granules, or with each other (**Fig. 1d, Extended Data Fig. 3**). This result indicates that the *TCRA* and *TCRB* mRNAs are not translationally repressed but may be localized to translation “hot spots.”

To capture actively translated *TCRA* and *TCRB* mRNAs and examine their interactions with eIF3, we analyzed polysomes in activated Jurkat cells (**Fig. 1e, Extended Data Fig. 4**). We compared these mRNAs to another pan-crosslinked mRNA, *DUSP2*, and to two mRNAs that crosslinked to eIF3 only through their 5’-UTRs (*EGR1, TRIM28*). To stabilize eIF3 interactions, we first treated activated Jurkat cells with the protein-protein crosslinker dithiobis(succinimidyl propionate) (DSP). We then incubated the cell lysates with RNaseH and DNA oligonucleotides designed to cleave the mRNAs between the 5’-UTR, coding sequence (CDS), and 3’-UTR, and isolated the polysomes on sucrose gradients (**Fig. 1e-f**) (**Supplementary Table 7**). This protocol efficiently cleaved the mRNAs, and prevented the released 5’-UTR and 3’-UTR elements from entering polysomes (**Extended data Fig. 4b-e**). This allowed us to detect eIF3 interactions with the mRNA CDS regions independent of eIF3 interactions with the UTR sequences identified in the PAR-CLIP analysis. To detect mRNAs interacting with eIF3 in the polysomes we performed anti-EIF3B immunoprecipitations followed by qRT-PCR (**Fig. 1e**). Using primers to the CDS regions of the mRNAs, we found that eIF3 only immunoprecipitated the pan-crosslinked mRNAs (*TCRA, TCRB, DUSP2*) from polysomes, but not mRNAs that only crosslinked to eIF3 through their 5’-UTRs (*EGR1, TRIM28*) (**Fig. 1g-h**). Importantly, all of these mRNAs are present in translating ribosomes and can be immunoprecipitated with eIF3 when the mRNAs are left intact (RNaseH treatment without DNA oligos) (**Fig. 1g**). These results indicate that eIF3 remains bound to the coding sequences (CDS) of the pan-mRNAs *TCRA, TCRB* and *DUSP2* independent of their 5’-UTR and 3’-UTR elements. By contrast mRNAs that crosslink to eIF3 only through their 5’-UTRs do not remain bound to eIF3 in polysomes. These results further support the model that *TCRA* and *TCRB* mRNA puncta represent translation “hot spots,’’ and the pan-mRNA crosslinking pattern is a result of eIF3 remaining engaged with these mRNAs during translation elongation.

We next tested whether eIF3 interactions with the *TCRA* and *TCRB* mRNAs encoding the major components of the TCR play a direct role in T cell activation, as the pan-mRNA crosslinking pattern to these mRNAs only occurs in activated Jurkat cells. We first probed the role of eIF3 interactions with the *TCRA* and *TCRB* mRNA 3’-UTRs, since eIF3 interactions with cellular mRNA 3’-UTRs have not been described before. We used CRISPR-Cas9 genome editing to delete the entire PAR-CLIP site in each of the 3’-UTRs of *TCRA* and *TCRB* mRNA, generating *TCRA* Δ*PAR* and *TCRB* Δ*PAR* Jurkat cells (**Extended Data Fig. 5a, Supplementary Table 7**). We then activated the cells with anti-CD3 and anti-CD28 antibodies, which induce both TCR and CD28 coreceptor signaling required for full T cell activation^15^. We analyzed the expression of total TCR protein levels in the *TCRA* Δ*PAR* and *TCRB* Δ*PAR* cells compared to WT Jurkat cells using western blots and an anti-TCRA antibody, since the formation of an intact TCR is required to stabilize both the TCRA and TCRB subunits^16,17^. In the anti-CD3/anti-CD28 activated WT Jurkat cells, TCR expression occurred as a burst that peaked 5-8 hours after stimulation (**Fig. 2a**). By contrast, both *TCRA* Δ*PAR* and *TCRB* Δ*PAR* Jurkat cell populations expressed lower levels of the TCR proteins compared to WT cells, and failed to show a burst in TCR expression at early time points after activation (**Fig. 2a**).

**Figure 2.**
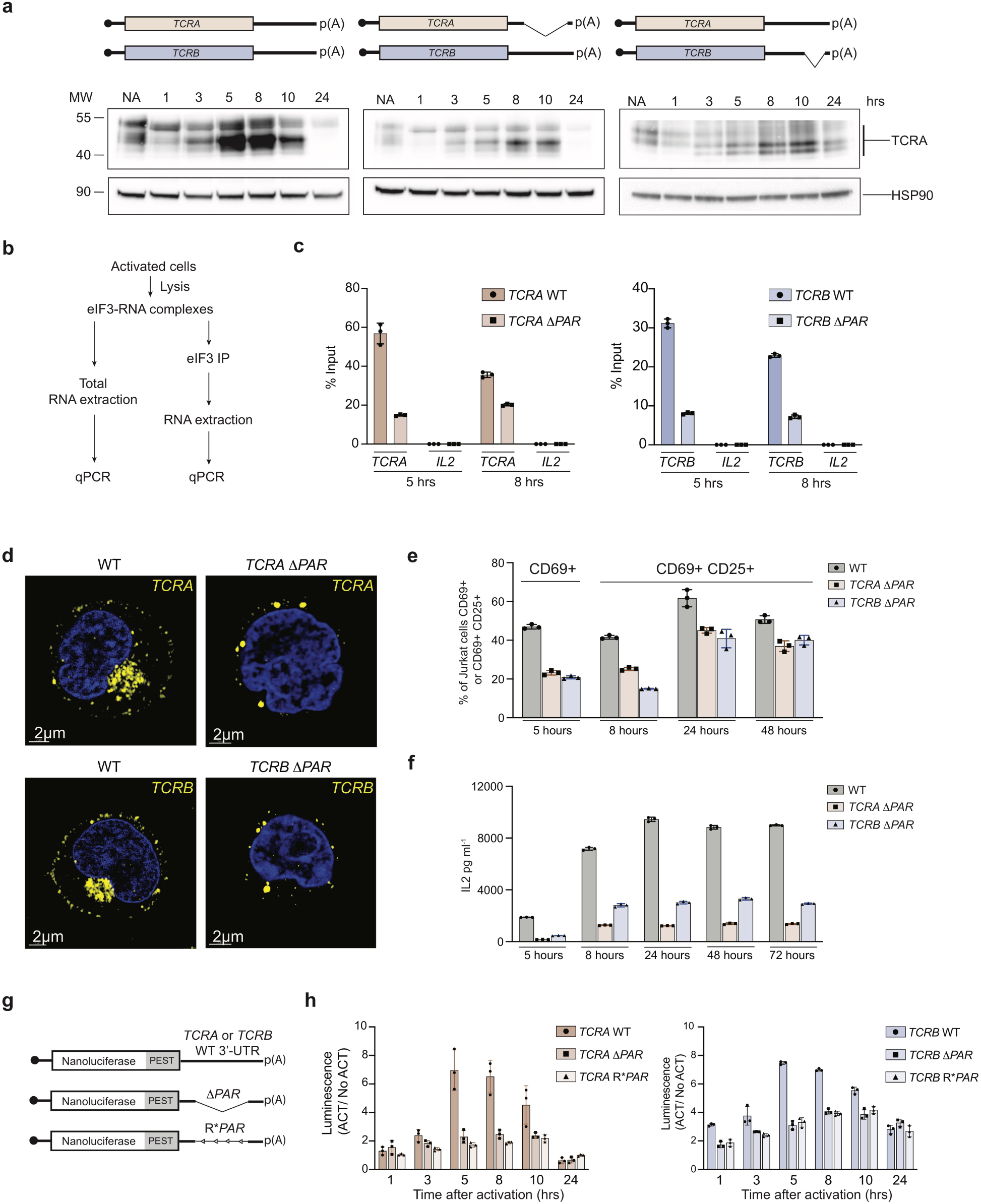
Role of *TCRA* and *TCRB* mRNA 3’-UTR elements in TCR expression and localization in activated Jurkat cells. **a**, Western blots of TCRA protein levels as a function of time after anti-CD3/anti-CD28 activation. Cell lines used: WT, wild-type Jurkat cells, *TCRA* Δ*PAR* and *TCRB* Δ*PAR*, Jurkat cells in which the eIF3 PAR-CLIP sites in the 3’-UTRs of the *TCRA* and *TCRB* mRNAs, respectively, have been deleted. Schematics of *TCRA* and *TCRB* mRNAs with and without eIF3 PAR-CLIP sites are shown above. HSP90 was used as a loading control. **b**, Schematic of immunoprecipitation of eIF3 using an anti-EIF3B antibody^1^, followed by qRT-PCR to quantify the amount of *TCRA, TCRB* or *IL2* mRNA bound to eIF3. **c**, Immunoprecipitation of eIF3 using an anti-EIF3B antibody^1^, followed by qRT-PCR to quantify the amount of *TCRA* mRNA (left) or *TCRB* mRNA (right) bound to eIF3. Jurkat cells with or without the eIF3 PAR-CLIP site in the 3’-UTR were analyzed after activation with anti-CD3/anti-CD28 antibodies for 5 hours or 8 hours. The percent mRNA bound to anti-EIF3B beads is calculated relative to total mRNA isolated from the cells. (*n* = 3, with mean and standard deviations shown). **d**, Measurement of TCR cluster formation in WT, *TCRA* Δ*PAR* and *TCRB* Δ*PAR* Jurkat cells, after 5 hours of activation using anti-TCRA/anti-TCRB protein staining and Airyscan confocal microscopy. (*n = 5* for each time point after activation for all cell lines, with representative images shown.) **e**, Flow cytometric analysis of cells expressing T cell activation markers CD69 (early activation marker) and CD25 (mid-activation marker) in WT, *TCRA* Δ*PAR* or *TCRB* Δ*PAR* Jurkat cells at different time points after activation with anti-CD3/anti-CD28 antibodies. (*n* = 3, with mean and standard deviations shown). P < 0.0001 for CD69 expression in *TCRA* Δ*PAR* and *TCRB* Δ*PAR* cells relative to WT cells at 5 hours, using one-way ANOVA. P < 0.001 for CD69 plus CD25 expression in *TCRA* Δ*PAR* and *TCRB* Δ*PAR* cells relative to WT using two-way ANOVA. **f**, Quantification of secreted IL2 from WT, *TCRA* Δ*PAR* or *TCRB* Δ*PAR* cells at different time points after stimulation with anti-CD3/anti-CD28 antibodies, as determined by ELISA. (*n* = 3, with mean and standard deviation shown.) P < 0.0001 for *TCRA* Δ*PAR* and *TCRB* Δ*PAR* cells relative to WT cells, as determined by two-way ANOVA. **g**, Schematic of nanoluciferase reporters stably expressed in Jurkat cells. The reporters carry the *HBB* 5’-UTR and WT, *ΔPAR* or R\**PAR* 3’-UTRs. WT, intact 3’-UTR from either *TCRA* or *TCRB* mRNA; *ΔPAR*, 3’-UTR of *TCRA* or *TCRB* with the eIF3 PAR-CLIP site deleted; R\**PAR*, reversed PAR-CLIP sequence in the 3’-UTR of *TCRA* or *TCRB* mRNA. **h**, Luciferase activity in anti-CD3/anti-CD28 activated Jurkat cells, relative to non-activated controls (*n* = 3, with mean and standard deviations shown). P < 0.0001 at 5 hr and 8 hr, and P ≤ 0.0002 for 1hr, for *TCRA* Δ*PAR* and R\**PAR* relative to WT, using two-way ANOVA. P < 0.0001 for *TCRB* Δ*PAR* and R\**PAR* relative to WT for 1 hr - 10 hr, using two-way ANOVA.

Importantly, *TCRA* and *TCRB* mRNA levels were unaffected or even increased in the *TCRA* Δ*PAR* and *TCRB* Δ*PAR* Jurkat cells (**Extended Data Fig. 5b, c**) suggesting that TCR expression levels are regulated post-transcriptionally, and disrupting eIF3 interactions with the *TCRA* or *TCRB* mRNA 3’-UTR eliminates the early burst in TCR expression. To test for eIF3 binding, we used anti-EIF3B immunoprecipitation followed by qRT-PCR to measure the amount of *TCRA* and *TCRB* mRNAs that interact with eIF3 in the wild-type (WT), *TCRA* Δ*PAR*, and *TCRB* Δ*PAR* cells after anti-CD3/anti-CD28 activation (**Fig. 2b**). The amount of *TCRA* and *TCRB* mRNAs bound to eIF3 was significantly lower in the *TCRA* Δ*PAR* and *TCRB* Δ*PAR* cells compared to WT cells at both the 5 hour and 8 hour timepoints (**Fig. 2c**), indicating that deleting the eIF3 PAR-CLIP sites in the 3’-UTR disrupts eIF3 interactions with these mRNAs substantially. Taken together, these results support the model that eIF3 binding to the 3’-UTR elements of the *TCRA* and *TCRB* mRNAs mediates the early burst in TCR translation after activation.

Since *TCRA* Δ*PAR* and *TCRB* Δ*PAR* Jurkat cells did not show the transient burst in TCR expression after activation, we tested whether deletion of the eIF3 PAR-CLIP sites in the 3’-UTRs of *TCRA* and *TCRB* mRNAs caused a general deficiency in T cell activation. We first tested whether the reduced amount of TCR expression levels observed in *TCRA ΔPAR* and *TCRB ΔPAR* cells at the earlier time points after activation affected TCR cluster formation, a central step in immune synapse formation^18^. We performed immunofluorescence on WT, *TCRA ΔPAR* and *TCRB ΔPAR* Jurkat cells using anti-TCRA and anti-TCRB antibodies to detect the TCR. In contrast to WT cells, both *TCRA ΔPAR* and *TCRB ΔPAR* cells failed to form TCR clusters by 5 hours after stimulation (**Fig. 2d**). Both mutants did form TCR clusters at 8 hours after stimulation, when TCR expression is slightly increased in these cells (**Extended Data Fig. 5d**), indicating that TCR protein expression levels and TCR cluster formation are tightly coupled. We then measured the T cell activation markers CD69 and CD25, expressed at early-to-mid timepoints after activation, by flow cytometry. Both *TCRA* Δ*PAR* and *TCRB* Δ*PAR* Jurkat cells had lower cell populations expressing CD69 and CD25 compared to WT cells (**Fig. 2e, Extended Data Fig. 5e**). We also measured the cytokine IL2, which is secreted at later time points after activation. Both *TCRA* Δ*PAR* and *TCRB* Δ*PAR* Jurkat cells secreted lower amounts of IL2 compared to WT cells (**Fig. 2f**). Taken together, these results show that deletion of the eIF3 PAR-CLIP site in either the *TCRA* or *TCRB* 3’-UTR results in decreased eIF3 interactions with these mRNAs and eliminates the burst in TCR expression, which cascades into multiple defects in early and late steps in T cell activation, from TCR clustering to IL2 secretion.

To further define the functional role of eIF3 interactions with the 3’-UTRs of the *TCRA* and *TCRB* mRNAs, we constructed nanoluciferase reporters fused to the WT *TCRA* or *TCRB* mRNA 3’-UTR sequence, or 3’-UTRs with the eIF3 PAR-CLIP site deleted (*ΔPAR*), or 3’-UTRs with the reversed sequence of the eIF3 PAR-CLIP site (R\**PAR*, i.e. 5’-3’ sequence reversed to 3’-5’ direction) (**Fig. 2g**). Jurkat cells expressing the reporters with the WT *TCRA* or *TCRB* mRNA 3’-UTR sequences had higher luminescence that peaked 5-8 hours after activation, while cells expressing nanoluciferase from reporters with a deletion or reversal of the eIF3 PAR-CLIP site sequence showed no apparent translation burst (**Fig. 2h**). These results recapitulate the 3’-UTR dependence of the translational burst for the endogenous *TCRA* and *TCRB* mRNAs (**Fig. 2a**). Also as observed for the *TCRA* Δ*PAR* and *TCRB* Δ*PAR* Jurkat cell lines, cells expressing reporters with either *TCRA* Δ*PAR* or R\**PAR* or *TCRB* Δ*PAR* or R\**PAR* 3’-UTRs had no significant defect in the nanoluciferase mRNA levels compared to reporters with the corresponding WT 3’-UTR sequences (**Extended Data Fig. 5f**). Finally, immunoprecipitation of eIF3 followed by qRT-PCR quantification of nanoluciferase mRNA showed that less nanoluciferase mRNA was bound to eIF3 when the 3’-UTR PAR-CLIP site was either deleted or reversed, compared to nanoluciferase mRNAs carrying the WT *TCRA* or *TCRB* 3’-UTR (**Extended Data Fig. 5g**). Taken together, the experiments with nanoluciferase reporters show that the *TCRA* and *TCRB* 3’-UTRs are necessary and sufficient to drive the translational burst seen after Jurkat cell activation, and correlates with eIF3 binding to the eIF3 PAR-CLIP sequences.

The eIF3-mediated burst in TCR expression occurred when Jurkat cells were fully activated through TCR and CD28 signaling via anti-CD3 and anti-CD28 antibodies. Interestingly, the CD28 costimulatory signal alone was sufficient to cause the transient burst in TCRA protein levels in the early time points after activation (**Fig. 3a**), indicating that the CD28 costimulatory pathway drives the transient burst in TCR protein expression in Jurkat cells. CD28 signaling involves multiple membrane-associated events^19^. Notably, anti-CD28 stimulation was sufficient to induce a transient burst in nanoluciferase expression from reporters harboring the WT *TCRA* mRNA 3’-UTR, but only when the reporters were tethered to the membrane via a CD3zeta-derived N-terminal sequence (**Fig. 3b-e**). This burst did not occur when reporters harboring the *TCRA* 3’-UTR lacked the eIF3 PAR-CLIP site (*TCRA* Δ*PAR*) (**Fig. 3c,d**). Furthermore, activation using only anti-CD3 antibodies led to increased reporter expression, but the increase peaked at a later time and did not drop off as significantly at later time points (**Fig. 3f**). These data indicate that the transient burst in TCR expression likely requires not only specific interactions between eIF3 and the *TCRA* and *TCRB* 3’-UTRs but also membrane-proximal CD28 signaling, as CD3zeta is co-translationally inserted into the membrane^20^.

**Figure 3.**
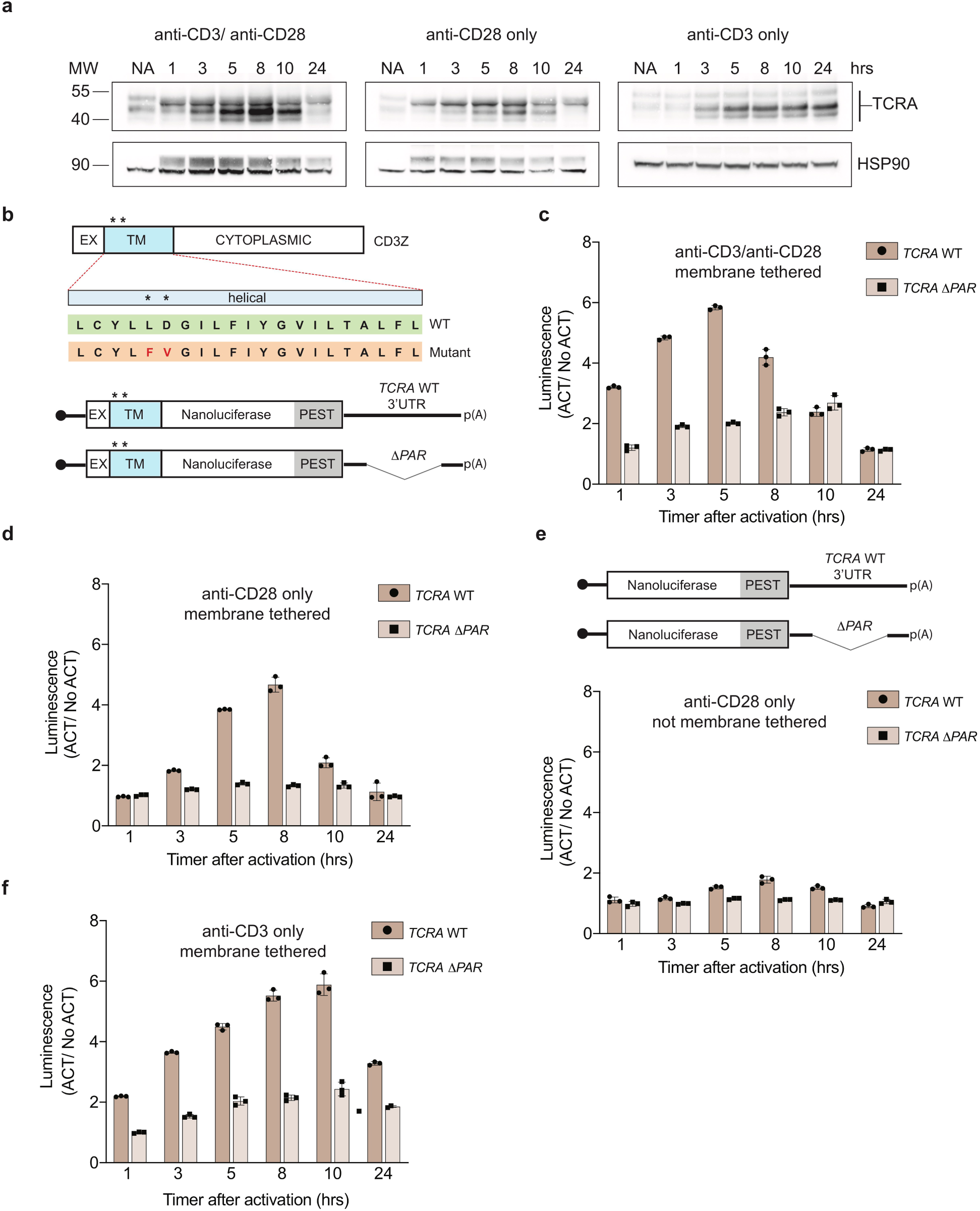
CD28-mediated burst in TCR and nanoluciferase reporter translation mediated by *TCRA* and 3’-UTRs. **a**, Western blots of TCRA protein levels in Jurkat cells as a function of time after different modes of activation. HSP90 was used as a loading control. **b**, Top, schematic of wild-type CD3-zeta protein with asterisk indicating the two amino acids mutated in the TM region (EX: extracellular, TM: transmembrane). Bottom: Schematic of the membrane-tethered nanoluciferase reporter stably expressed in Jurkat cells. Nanoluciferase is fused C-terminal to the extracellular and transmembrane segments of CD3-zeta, mutated to prevent TCR association. The reporters carry the *HBB* 5’-UTR and WT *TCRA* or *TCRA* Δ*PAR* 3’-UTR. **c**, Luciferase activity in Jurkat cells expressing membrane-tethered reporters stimulated with anti-CD3/anti-CD28 antibodies, relative to non-activated controls (*n* = 3, with mean and standard deviations shown). P < 0.0001 for *TCRA* Δ*PAR* relative to WT from 1-8 hr, and P ≤ 0.04 for 10 hr, using two-way ANOVA. **d**, Luciferase activity in Jurkat cells activated with anti-CD28 only, relative to non-activated controls (*n* = 3, with mean and standard deviations shown). P < 0.0001 for *TCRA* Δ*PAR* relative to WT from 3 hr to 10 hr, using two-way ANOVA. **e**, Luciferase activity in Jurkat cells expressing non-tethered nanoluciferase reporters (schematic at the top) and stimulated with only anti-CD28 antibodies. Values are normalized to non-activated controls (*n* = 3, with mean and standard deviations shown). P <0.001 for *TCRA* Δ*PAR* relative to WT at 5 hr to 10 hr, using two-way ANOVA. **f**, Luciferase activity in Jurkat cells expressing the tethered reporters in **b**, and stimulated with anti-CD3 antibodies, relative to non-activated controls (*n* = 3, with mean and standard deviations shown). P < 0.001 for *TCRA* Δ*PAR* relative to WT at 1hr to 24 hrs, using two-way ANOVA.

Jurkat cells may not fully recapitulate the events that occur early in the activation of primary T cells. We therefore tested the importance of eIF3 interactions with the *TCRA* and *TCRB* mRNA 3’-UTRs in primary T cells. As in the Jurkat cells, we used CRISPR-Cas9 genome editing to delete the eIF3 PAR-CLIP sites in each of the *TCRA* and *TCRB* mRNA 3’-UTRs in primary human T cells from two healthy human donors (**Extended Data Fig. 5a, Supplementary Table 7**). PCR analysis showed successful deletion of the eIF3 PAR-CLIP site in the *TCRA* 3’-UTR or in the *TCRB* 3’-UTR (*TCRA* Δ*PAR* or *TCRB* Δ*PAR*, respectively) in approximately 43% - 49% of the cells (**Extended Data Fig. 6a**). For comparison with the *TCRA* Δ*PAR* or *TCRB* Δ*PAR* cell populations, we used cells edited with a scrambled sgRNA (SC), which does not target any site in the human genome. We first measured total TCR protein levels by western blot, after different times following T cell activation using anti-CD3 and anti-CD28 antibodies (**Fig. 4a**). The SC control cells – which should behave as wild-type T cells – exhibited a substantial burst in TCRA protein levels immediately after activation (∼1 hr). By contrast, in both *TCRA* Δ*PAR* and *TCRB* Δ*PAR* cell populations, TCRA protein levels were nearly absent or clearly reduced at early time points after activation (**Fig. 4a**). Only at later time points did TCRA levels in the *TCRA* Δ*PAR* and *TCRB* Δ*PAR* cell populations begin to increase. The early burst in TCRA translation depended on anti-CD28 stimulation, as observed in Jurkat cells (**Extended Data Fig. 6b**). Next we checked whether the reduced TCR protein levels in *TCRA* Δ*PAR* and *TCRB* Δ*PAR* cell populations at early time points after activation affected the rate of TCR clustering. We used the same cell populations as those for western blot analysis (**Fig. 4a**), to correlate the TCRA protein levels observed in the western blots with TCR cluster formation. Using TCR protein staining, we observed a decrease in the proportion of cells that formed TCR clusters in the *TCRA* Δ*PAR* and *TCRB* Δ*PAR* cell populations compared to SC control cells, with more severe defects seen at early time points (**Fig. 4b**). We also stained cells with a fluorescently-tagged anti-TCR antibody and used flow cytometric analysis to measure the levels of TCR on the cell surface with and without activation with I+PMA (**Extended Data Fig. 6c,d**). A substantially lower percentage of the *TCRA* Δ*PAR* and *TCRB* Δ*PAR* cell populations expressed TCR on the cell surface compared to SC control cells or cells edited with a single gRNA (**Extended Data Fig. 6c,d**). Taken together, these results recapitulate those seen in Jurkat cells, but on a shorter time frame, with the burst in TCR expression occurring ∼1 hour in primary T cells, rather than ∼5 hours in Jurkat cells.

**Figure 4.**
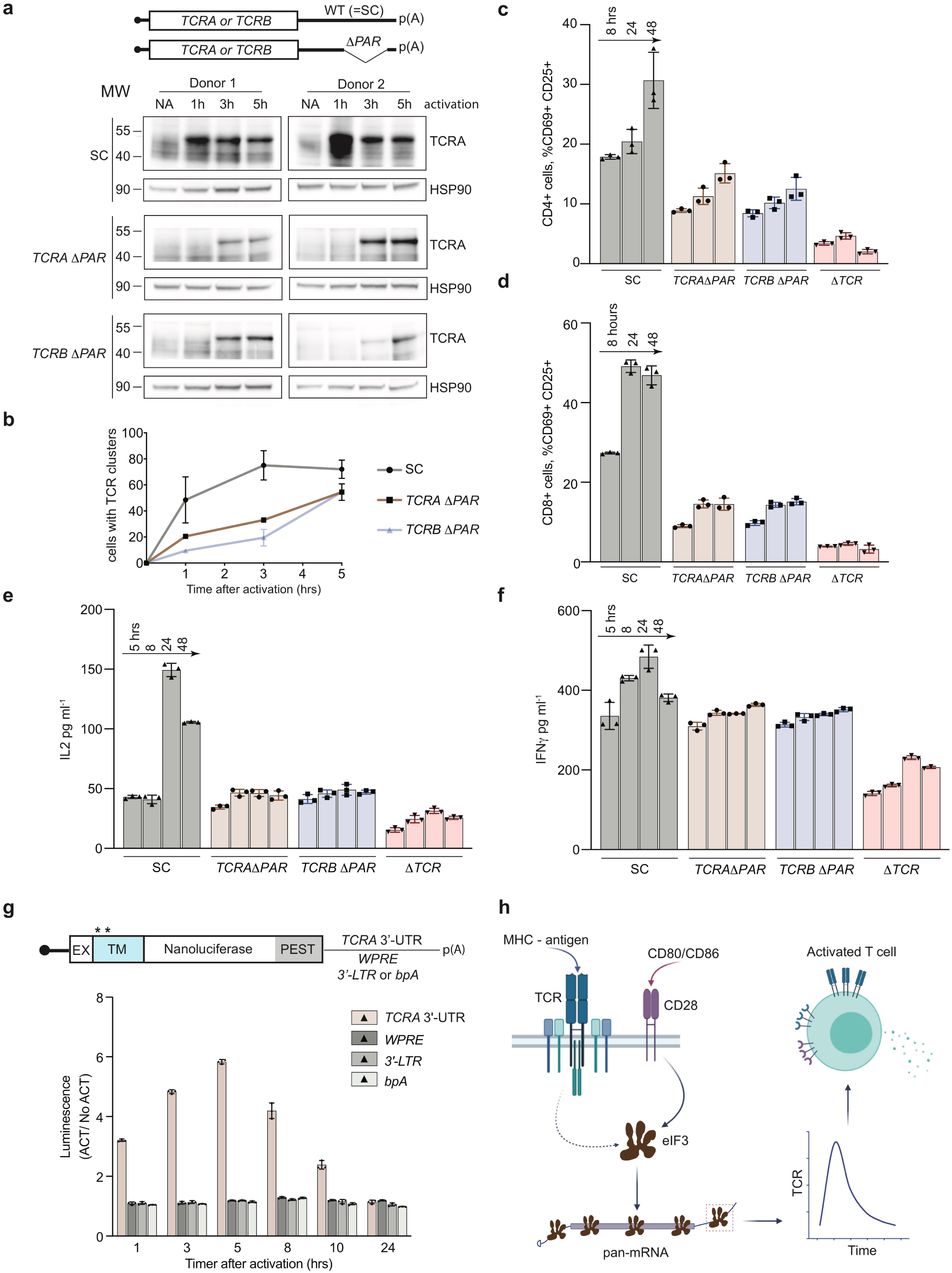
Role of the *TCRA* and *TCRB* mRNA 3’-UTR elements in primary human T cells. **a**, Western blots measuring TCRA protein levels as a function of time after anti-CD3/anti-CD28 activation. Cell lines used are labeled on the left: *TCRA* Δ*PAR, TCRB* Δ*PAR*, and SC (scrambled gRNA). HSP90 was used as a loading control. Schematics of *TCRA* and *TCRB* mRNAs with and without eIF3 PAR-CLIP sites are shown above. SC control cells have the wild-type (WT) 3’-UTRs for *TCRA* and *TCRB* mRNAs. **b**, The number of T cells with one or more TCR clusters measured by anti-TCRA/anti-TCRB protein staining and epifluorescence microscopy as a function of time after anti-CD3/anti-CD28 activation. A total of 100 cells from each donor were imaged for *TCRA* Δ*PAR* (*n* = 2 donors, stained with anti-TCRA antibody), *TCRB* Δ*PAR* (*n* = 2 donors, stained with anti-TCRB antibody), and SC cell lines (*n* = 2 donors, each stained separately with anti-TCRA and anti-TCRB antibodies). Values are mean ± standard deviation. P < 0.005 comparing *TCRA* Δ*PAR* and *TCRB* Δ*PAR* to SC after 1 hour of activation. P ≤ 0.0002 comparing *TCRA* Δ*PAR* and *TCRB* Δ*PAR* to SC after 3 hours of activation, using two-way ANOVA. **c**, Flow cytometric analysis measuring T cell activation markers CD69 (early activation marker) and CD25 (mid-activation marker), quantifying the mean percent of CD4+ T cells that are CD69+ CD25+ from (*n* = 2 donors). **d**, Flow cytometric analysis of CD8+ T cells, quantifying the mean percent of CD8+ T cells that are CD69+ CD25+ from (*n* = 2 donors). Cells sorted as shown in **Extended Data Fig. 7. e-f**, Quantification of **e**, secreted IL2 and **f**, secreted IFNγ from SC, *TCRA* Δ*PAR, TCRB* Δ*PAR* and Δ*TCR* cell populations from (*n* = 2 donors) at different time points after stimulation with anti-CD3/anti-CD28 antibodies, as determined by ELISA. Values are mean ± standard deviation, P < 0.0001 for *TCRA* Δ*PAR* and *TCRB* Δ*PAR* relative to SC at 24 hrs and 48 hrs for secreted IL2, and P < 0.0001 for *TCRA* Δ*PAR* and *TCRB* Δ*PAR* relative to SC at 8 hrs and 24 hrs for secreted IFNγ, using two-way ANOVA. **g**, Luciferase activity in Jurkat cells stably expressing membrane-tethered nanoluciferase reporters activated with anti-CD3/anti-CD28 antibodies, as diagrammed at the top, normalized to luciferase activity in non-activated Jurkat cells. (*n* = 3, with mean and standard deviations shown). The nanoluciferase reporters have a *HBB* 5’-UTR and either the intact 3’-UTR from *TCRA*, the Woodchuck Hepatitis Virus Posttranscriptional Regulatory Element (*WPRE*), the gammaretroviral 3’-Long terminal repeat (*3’-LTR*), or the bovine growth hormone polyadenylation signal (*bpA*) following the nanoluciferase coding sequence. P < 0.0001 for *WPRE, 3’-LTR*, or *bpA* relative to *TCRA* 3’-UTR at 5 hrs - 10 hrs, using two-way ANOVA. Luciferase data for the *TCRA* 3’-UTR reporter is shown from **Fig. 3c** for reference. The experiments were carried out at the same time. **h**, Model for early events in T cell activation, where eIF3 interactions with TCR mRNA 3’-UTR sequences lead to a burst in TCR translation and downstream T cell function. eIF3 structure from^48^ and figure partially created in BioRender.

To test whether the defect in TCR clustering in the *TCRA* Δ*PAR* and *TCRB* Δ*PAR* cell populations reflects a general deficiency in primary T cell activation, as observed in Jurkat cells, we measured the T cell activation markers CD69 and CD25 by flow cytometry and secreted cytokines IL2 and IFNγ using ELISA (**Fig. 4c, d, Extended Data Fig. 7**). Fewer cells in the *TCRA* Δ*PAR* and *TCRB* Δ*PAR* CD4+ and CD8+ primary cell populations expressed CD69 at early time points after activation (5-8 hours) and fewer expressed both CD69 and CD25 at later time points after activation, compared to SC control cells (**Fig. 4c, d, Extended Data Fig. 8**). Furthermore, the *TCRA* Δ*PAR* and *TCRB* Δ*PAR* T cell populations secreted lower amounts of cytokines IL2 and IFNγ, compared to the SC control cells (**Fig. 4e, f, Extended Data Fig. 9**). Taken together, the *TCRA* Δ*PAR* and *TCRB* Δ*PAR* primary T cell populations exhibited multiple early and late activation defects, recapitulating the results obtained in Jurkat cells. These results further support the model that after T cell activation, eIF3 binding to the *TCRA* and *TCRB* mRNA 3’-UTRs drives an early burst in TCR translation that is required for subsequent steps in T cell activation.

T cells engineered to express chimeric antigen receptors (CARs) for cancer immunotherapy now in use clinically^21,22,23^ and in next-generation designs^24^ express the CAR from stably integrated transgenes that employ artificial 3’-UTRs in the CAR-encoding mRNA. To test whether these 3’-UTRs induce a transient burst in translation as observed with the WT *TCRA* or *TCRB* 3’-UTRs, we fused 3’-UTR sequences now in use for CAR expression to nanoluciferase reporters and expressed these in Jurkat cells (**Fig. 4g**). In contrast to the *TCRA* 3’-UTR (**Fig. 3**), the other 3’-UTR elements failed to induce the burst in nanoluciferase expression (**Fig. 4g**). These data suggest that fusing the *TCRA* or *TCRB* 3’-UTR sequences to engineered CAR genes could be used to obtain a more physiological expression pattern seen for the endogenous TCR.

Recent experiments indicate that T cells must cross a “threshold” of T cell receptor signaling to commit to activation and proliferation^25,26^, setting up a “digital” response to antigen recognition^25–28^. The response threshold involves integration of intensity and duration of TCR signaling^25,26,28^, and spans a wide range of TCR antigen affinity^25–28^. Notably, T cell commitment to clonal expansion and differentiation can occur with as little as 1 to 2 hours of TCR stimulation for effector CD4+ and naive CD8+ T cells^29,30^. This time period spans the burst in TCR protein synthesis mediated by eIF3 interactions with the *TCRA* and *TCRB* 3’-UTR elements (**Fig. 4a**). Naive CD4+ T cells require a longer duration of TCR signaling of ∼20 hours^29,31^. Although we were not able to distinguish levels of TCR translation in isolated CD8+ and CD4+ T cells (**Fig. 4**), subsequent events in T cell activation including CD69 and CD25 expression, and IL2 and IFNγ secretion, were equally affected in CD8+ and CD4+ cells in which the 3’-UTR PAR-CLIP sites were deleted (**Fig. 4c-f**). In an immune response, CD28 engagement serves as the second signal required for T cell activation^15,32^ and affects the first minutes of TCR-mediated signaling^33–37^. Here we now show CD28-mediated signaling is also needed for the burst of TCR translation on the hour timescale (**Fig. 3, Extended Data Fig. 6b**). Taken together, our results indicate that eIF3 controls *TCRA* and *TCRB* mRNA translation during the first critical hours after antigen recognition that lead to subsequent T cell commitment to proliferation and differentiation (**Fig. 4h**). Given that CD28 is required for PD-1 mediated inhibition of T cell activation^38,39^, it will be interesting to determine the role of eIF3-mediated control of TCR expression in PD-1 checkpoint blockade-based cancer immunotherapy^40^. Additionally, our results using nanoluciferase reporters suggest that eIF3-responsive mRNA 3’-UTR elements could be used to improve chimeric antigen receptor expression and CAR-T cell responsiveness^24,41^.

The eIF3 PAR-CLIP experiment we present here provides a snapshot of eIF3-RNA interactions that occur at the time TCR translation is most sensitive to eIF3 regulation (5 hours in Jurkat cells, **Fig. 2a**). At this point in time, eIF3 crosslinks to multiple mRNAs encoding proteins involved in immune cell function (**Supplementary Table 6**). Interestingly, the patterns of eIF3 crosslinking, which for a number of mRNAs include interactions with the protein coding sequence and 3’-UTR (**Fig. 1, Extended Data Fig. 2e**), suggest an active role for eIF3 in promoting translation of these mRNAs. In support of this model, the two examples of pan-mRNAs we examined here (*TCRA* and *TCRB*) reside in puncta distinct from P bodies and stress granules (**Fig. 1d**), and bind to eIF3 via the mRNA coding sequence in translating ribosomes (**Fig. 1e-h**). However, this eIF3-mediated translation activation is transient, lasting only 1-2 hours in primary T cells (**Fig. 4a, Extended Data Fig. 6b**). Recent evidence suggests that eIF3 can remain associated with translating ribosomes^42–44^, a phenomenon that seems to be enhanced for the pan-mRNAs identified here. Notably, additional layers of translation regulation also contribute to T cell function^45^, particularly with respect to mTOR signaling^10,46^ and carbon metabolism^11,47^. The present PAR-CLIP experiments should help to elucidate additional roles for eIF3-mediated translation regulation and to more fully map the system-wide role of translation regulation in T cell activation.

## Materials and methods

### Jurkat cell culture

Human Jurkat, Clone E6-1 (ATCC TIB-152) was purchased from American Type Culture Collection (ATCC) and was maintained in RPMI 1640 Medium (ATCC modification) with 10% FBS (VWR Life Science Seradigm) and 0.01% Penicillin-Streptomycin (10,000 U/mL). The cells were maintained between 1 million to 8 million cells/mL. When cells were stimulated they were always maintained at 8 million cells/mL.

### Isolation of human primary T cells

Primary human T cells were isolated from healthy human donors from leukoreduction chambers after Trima Apheresis (Vitalant, formerly Blood Centers of the Pacific). Peripheral blood mononuclear cells (PBMCs) were isolated from whole blood samples by Ficoll centrifugation using SepMate tubes (STEMCELL) per manufacturer’s instructions. T cells were isolated from PBMCs from all cell sources by magnetic negative selection using an EasySep Human T Cell Isolation Kit (STEMCELL), per manufacturer’s instructions. Unless otherwise noted, isolated T cells were stimulated as described below and used immediately.

### Primary human T cell culture

Bulk T cells were cultured in XVivo15 medium (STEMCELL) with 5% fetal bovine serum (FBS), 50 μM 2-mercaptoethanol, and 10 μM *N*-acetyl-cystine. Immediately after isolation, T cells were stimulated for 2 days with anti-human CD3/CD28 magnetic Dynabeads (ThermoFisher) at a bead to cell concentration of 1:1, along with cytokine IL-2 at 500 U/mL (UCSF Pharmacy). After electroporation, T cells were cultured in media with IL-2 at 500 U/mL. Throughout the culture period T cells were maintained at an approximate density of 1 million cells per mL of media. Every 2–3 days after electroporation, additional media was added, along with additional fresh IL-2 to bring the final concentration to 500 U/mL, and the cells were transferred to larger culture flasks as necessary to maintain a density of 1 million cells per mL.

For the T cell stimulation assay, edited primary T cells were transferred to fresh media lacking IL2 after 9 days of culturing. The T cells were then stimulated with anti-CD3 and anti-CD28 antibodies using flat bottom plates coated with anti-CD3 antibody (at a 10 μg/mL concentration), and anti-CD28 antibody added to the cell culture media at a concentration of 5 µg/mL.

### Jurkat cell stimulation

The Jurkat cells used for the for the PAR-CLIP experiment were stimulated with 1X Cell Stimulation Cocktail, containing phorbol 12-myristate 13-acetate (PMA) and ionomycin (Thermofisher, Catalog number: 00-4970-93) to ensure a large proportion of the cells were activated. Unless otherwise stated, all other experiments involving activated Jurkat cells used anti-CD3 and anti-CD28 antibodies (Tonbo) for stimulation. Flat bottom plates were coated with anti-CD3 antibody (at a 10 μg/mL concentration), and anti-CD28 antibody was added to the cell culture media at a concentration of 5 µg/mL.

### 4-thiouridine optimization for PAR-CLIP experiments

We used Jurkat cells as a model for T cells, as PAR-CLIP experiments require a large number of cells labeled with 4-thiouridine at a non-toxic concentration^14^. Jurkat cells also have a defined T cell receptor and transcriptome, avoiding the donor-to-donor variability of primary T cells. Jurkat cells were seeded, so that they reached 8×10^5^ cells ml^-1^ seeding density on the day of the experiment. Varying concentrations of 4-thiouridine (s4U) were added to the cells (50 µM, 75 µM, 100 µM, or none as a negative control). The cells were then incubated for different time points: 8 hours, 10 hours, 12 hours, or 16 hours. After each incubation time cell viability was determined using the CellTiter-Glo assay (Promega), according to the manufacturer’s instructions. Concentrations at which the relative luminescence in the presence of s4U (luminescence of the s4U treated cells/luminescence of the untreated cells) exceeded 95% were considered non-toxic. Based on these measurements, we used 50 µM of 4-thiouridine for PAR-CLIP experiments.

### PAR-CLIP

Two biological replicates were used to perform PAR-CLIP analysis as described in^1^, with modifications for Jurkat cells. 50 µM of 4-thiouridine was determined as non-toxic to Jurkat cells over the time course of the PAR-CLIP experiments^14^. A total of 55 million Jurkat cells seeded at 8 million cells ml^-1^ was treated with 50 µM of 4-thiouridine (Sigma) for 7 hours, then stimulated with 1X Cell Stimulation Cocktail for 5 hours in media containing 50 µM of 4-thiouridine (**Fig. 1a**). The same number of cells were treated with 50 µM of 4-thiouridine (Sigma) for 12 hours without stimulation as a non-activated control. The cells were then crosslinked on ice with 365 nm UV irradiation at an energy dose of 0.2 J cm^-2^. The cells were pelleted by centrifugation at 100 x g for 15 min at 4 °C, and the pellet was resuspended in three volumes of NP40 lysis buffer (50 mM HEPES-KOH pH 7.5, 150 mM KCl, 2 mM EDTA, 0.5% Nonidet P-40 alternative, 0.5 mM dithiothreitol (DTT), 1 Complete Mini EDTA-free Protease Inhibitor Cocktail tablet (Roche)). The cell suspension was then incubated on ice for 10 min, passed through an 18G needle five times, and centrifuged at 13,000 x g for 15 min at 4 °C and RNAs were lightly digested by treatment with MNase (Thermo Scientific) at a final concentration of 0.05 U μl^-1^ for 20 min at 16 °C. For each PAR-CLIP assay 1000 μl of Dynabeads (Invitrogen) and 800 μl of anti-EIF3B antibody (Bethyl A301-761A) were used. The remaining steps of the PAR-CLIP analysis was performed exactly as described in ^1,50^ with the exception of using MNase at 5 U μl^-1^ for the on-bead digestion step.

### PAR-CLIP computational analysis

PAR-CLIP cDNA libraries were sequenced on an Illumina HiSeq 2500 platform. To eliminate potential PCR biases during PAR-CLIP library preparation, a random bar code was introduced into the 3’ adapter and all the reads that matched the random barcode were collapsed into single reads. Clusters of overlapping sequence reads mapped against the human genome version hg38 were generated using the PARalyzer software ^51^ incorporated into the PARpipe pipeline (https://ohlerlab.mdc-berlin.de/software/PARpipe_119/, ^13^) with the settings below. Binding sites were categorized using the Gencode GRCh38.p12 GTF annotations (gencode.v21.annotation.gtf), https://www.gencodegenes.org/human/.

The PARpipe settings used were

Conversion = T>C

Minimum read count per group = 5

Minimum read count per cluster = 7

Minimum read count for kde = 3

Minimum cluster size = 11

Minimum conversion locations for cluster = 2

Minimum conversion count for cluster = 2

Minimum read count for cluster inclusion = 1

Minimum read length = 20

Maximum number of non conversion mismatches = 1

### Comparison of eIF3 PAR-CLIP results in Jurkat and HEK293T cells

To compare RNAs in activated Jurkat cells crosslinked to eIF3 with those crosslinked to eIF3 in HEK293T cells^1^, the gene cluster lists (*.gene_cl.csv) from APA_REP1, APA_REP2, APB_REP1, APB_REP2, APD_REP1, and APD_REP2 were used (see **Supplementary Table 3**). The genes were first sorted by total read counts (“ReadCountSum”) from high to low for each library to obtain the top candidate genes. Then, the same number of top candidate genes from these sorted lists as the number of genes identified in HEK293T cells, for eIF3 subunit EIF3A, EIF3B, and EIF3D crosslinked to RNA^1^, were chosen for comparison.

### PAR-CLIP Pathway analysis

To determine biological pathways enriched in the set of mRNAs that crosslinked to eIF3 in activated Jurkat cells, genes with at least 100 total aligned reads were used, as determined in the PARpipe analysis described above^13^, from the EIF3A/C/B samples. Since PAR-CLIP reads are short, it is not possible to determine with certainty which mRNA transcript isoform cross-linked with eIF3. Therefore the most abundant mRNA transcript isoform for each gene was chosen, as determined by transcriptome profiling using kallisto (protein_coding category)^52^, as described in the Transcriptome Profiling section. Even with this choice, eIF3 crosslinks to mRNAs do not correlate with mRNA abundance (**Extended Data Fig. 2c**). Human genome GRCh38.p13 annotation was used to extract mRNA lengths by 5’-UTR, coding region and 3’-UTR (Ensembl Biomart)^53^. These genes were then sorted by the density of PAR-CLIP reads in the mRNA 5’-UTR region, prior to mapping pathways of transcripts that cross-linked to eIF3. Due to the complexity of *TCRA* and *TCRB* transcript annotation, these transcripts were excluded from the normalization calculation, but included in the pathway analysis. The top 500 genes from the resulting EIF3A/C/B PAR-CLIP gene lists were used, sorted as described above, to analyze gene enrichment profiles in the STRING database^54^. The top tissue-specific categories involve the immune system and T cell function (**Supplementary Table 6**). Note that the STRING database does not include TCR subunits in its analysis.

### Metagene analysis

The PAR-CLIP genes sorted as described above in the “PAR-CLIP pathway analysis” were used and mapped against the most abundant mRNA transcript isoforms to generate cumulative plots of the reads. Reads for the *TCRA* and *TCRB* mRNAs were manually extracted from the Bowtie version 1.0.0 Hg38 alignment of the eIF3 PAR-CLIP reads. We did not identify reads mapped to the D segment of *TCRB* (ie to *TRBD2*) due to its short length of 16 nucleotides. These reads were combined with the mapped reads in the *.read.csv files generated by Parpipe. The combined reads were then sorted to extract only reads annotated as 5’-UTR, start codon, coding, stop codon, and 3’-UTR. The most abundant transcript isoform, as identified in the Transcriptome Profiling section using kallisto (described below) was used. Reads mapped to the 5’-UTR and start codon were normalized by the length of the 5’-UTR. Reads mapped to the coding region and stop codon were normalized by the length of the coding region. Finally, reads mapped to the 3’-UTR were normalized to the length of the 3’-UTR. Relative positions of the mapped reads along a metagene were computed based on the locations each mapped read fell within its respective feature. Relative positions were from -1 to 0 for the 5’-UTR, 0 to 1 for the coding region, and 1 to 2 for the 3’-UTR. 5’-UTR values were computed by multiplying the relative position by -1, whereas the 3-UTR values were were computed by adding 1 to the relative position. Coding region relative positions were unchanged.

The empirical cumulative distribution frequency function from R package ggplot2^55^ was used to build the metagene plots from the vector of relative positions for the reads which mapped to a given set of reads. We defined mRNAs having a ratio of normalized 5’-UTR reads divided by 3’-UTR reads of 20 or more as “5’-UTR enriched” with respect to eIF3 crosslinking. All others were categorized as “pan-mRNAs,” with eIF3 crosslinking across the entire length of the mRNA. The cut-off value 20 is not a sharp boundary between the two categories of mRNA (**Extended Data Fig. 2d**).

### Transcriptome Profiling

RNA samples were extracted from HEK293T cells, non-activated Jurkat cells, and Jurkat cells activated for 5 hr with I+PMA, using the Direct-zol RNA Miniprep kit (Zymoresearch). The libraries were prepared using TruSeq Stranded RNA LT Ribo-Zero Gold kit (Illumina) following the manufacturer’s instructions, with two biological replicates per condition. Cutadapt (version 2.6)^56^ with a minimum read length of 20, 5’ end with a cutoff of 15 and the 3’ end with a cutoff of 10 in paired-end mode was used to remove adapters. RNA-seq reads were pseudoaligned using kallisto v.0.46.0 run in quant mode with default parameters to estimate transcript abundance (transcripts per million, TPM)^52^. The transcript index for kallisto was made with default parameters and GENCODE Release 32 (GRCh38.p13) FASTA file^57^.

To compare HEK293T or non-activated Jurkat mRNA abundance with that in activated Jurkat cells, the output from kallisto was used for analysis using DeSeq2 (ref 58). All protein-coding transcripts (protein_coding) were then selected for the comparisons, using tximport^59^ as part of the Bioconductor environment in R. All genes with ≥ 50 reads summed across all four samples in a comparison (two biological replicates per condition) were then used. Prior to making MA plots, the Approximate Posterior Estimation for generalized linear model (apeglm) shrinkage estimator for log fold changes was used60. DeSeq2 output in **Supplementary Table 5** is presented prior to shrinkage analysis.

### Polysome analysis of eIF3-associated mRNAs

Jurkat cells were seeded to reach 8 million cells ml^-1^ on the day of activation. Cells were stimulated with 1X Cell Stimulation Cocktail for 5 hours, and 5 minutes prior to cell harvest the cells were treated with 100 μg ml^-1^ cycloheximide. Cells were then collected into a 50 ml falcon tube and rinsed once with ice cold PBS supplemented with 100 μg ml^-1^ cycloheximide. The cells were then incubated with 0.5 mM of the the crosslinking reagent dithiobis(succinimidyl propionate) (DSP, Thermofisher) and 100 μg ml^-1^ cycloheximide in PBS at room temperature for 15 minutes, with rocking. The crosslinking reagent was then removed and the cells were incubated with quenching reagent (PBS, 100 μg ml^-1^ cycloheximide and 300 mM Glycine) for 5 minutes on ice. The cells were then rinsed again with ice cold PBS and flash frozen in liquid nitrogen.

A total of 4 x 10^8^ cells were lysed with 400 μl hypotonic lysis buffer (10 mM Hepes pH 7.9, 1.5 mM MgCl2, 10 mM KCl, 0.5 mM DTT, 1% Triton, 100 μg ml^-1^ cycloheximide, and one Complete EDTA-free Proteinase Inhibitor Cocktail tablet (Roche)). The cells were incubated for 10 min on ice and then passed through an 18G needle five times, and centrifuged at 13,000 × g for 15 min at 4 °C. The 400 μl supernatant was then transferred to a fresh eppendorf tube and subjected the RNaseH digestion by adding the following reagents: 3 mM MgCl2, 10 mM DTT, 200 units RNaseH (NEB) and 20 μl of SUPERasIN, with a total of 4 μM DNA oligos, as indicated in the figure legends. The mixture was then incubated at 37 °C for 20 minutes. After incubation 10 μl of the RNaseH treated lysate mixture was isolated to test the efficiency of the RNaseH digestion using qRT-PCR (**Extended Data Fig. 4d**) and the rest of the lysate was layered onto a 12 ml 10%-50% sucrose gradient, made with gradient buffer consisting of: 10% sucrose (w/v) or 50% sucrose (w/v), 100 mM KCl, 20 mM Hepes pH 7.6, 5 mM MgCl2, 1 mM DTT and 100 μg ml^-1^ cycloheximide. The gradient was centrifuged at 36,000 rpm (222k x g) for 2 hours at 4 °C in a SW-41 rotor. The gradient was then fractionated using the Brandel gradient fractionator and ISCO UA-6 UV detector and all the polysome fractions (∼ 5 ml) were collected into a fresh 15 ml falcon tube. 100 μl from each of the polysome fractions was kept aside to measure the input RNA amounts, and the rest of each polysome fraction was incubated with 100 μl of Dynabeads (Invitrogen) conjugated with 40 μl of anti-EIF3B antibody (Bethyl A301-761A) for 16 hours, rotating at 4 °C. After incubation, the beads were rinsed three times with 1000 μl room temperature NP40 lysis buffer (defined in the PAR-CLIP section), rotating for 5 minutes for each wash. After the final wash the beads were resuspended in 400 μl of Trizol (Thermofisher), the RNA was extracted and qRT-PCR was performed as described above.

### Mass spectrometry

To identify eIF3 subunits that crosslinked with RNAs in the PAR-CLIP experiments, a portion of eIF3 immunoprecipitated using Dynabeads as described above were treated with nonradioactive ATP during the T4 polynucleotide kinase labeling step. The nonradioactive samples were then run on the same gel next to the radiolabeled PAR-CLIP samples, Coomassie stained (Pierce) and the bands that matched with the phosphorimager printout were excised from the gel and submitted for identification using one-dimensional LC-MS/MS (**Supplementary Table 1**).

### sgRNA/Cas9 RNP production

The sgRNA/Cas9 RNPs used to edit Jurkat cells were produced by complexing sgRNA (Synthego) to Cas9 as described^61^ while RNPs to edit Primary Human T cells were produced by complexing a two-component gRNA (crRNA and tracrRNA, Dharmacon) to Cas9 as described in^62^. The targeting sequences for the sgRNAs and crRNA are given in **Supplementary Table 7**. Recombinant Cas9-NLS was obtained from MacroLab in the California Institute for Quantitative Biosciences.

### Jurkat and primary T cell genome editing

Jurkat cells used for electroporation were collected at passage 5 or lower and were maintained at a seeding density of 8 million cells ml^-1^ or lower. Primary T cells were isolated as described above. Prior to electroporation the Primary T cells were stimulated with magnetic anti-CD3/anti-CD28 Dynabeads (ThermoFisher) for 48 hours. After 48 hours these beads were removed from the cells by placing cells on an EasySep cell separation magnet for 2 min before electroporation. One million Jurkat (not activated) or primary T cells cells (activated with anti-CD3/anti-CD28 Dynabeads for 48 hours) were rinsed with PBS and then resuspended in 20 µl of Lonza electroporation buffer P3. The cells were then mixed with 2.5 μl Cas9 RNPs (50 pmol total) along with 2 μl of a 127-nucleotide non-specific single-stranded DNA oligonucleotide at 2 μg μl^-1^ (4 μg ssDNA oligonucleotide total). The cells were then electroporated per well using a Lonza 4D 96-well electroporation system with pulse code DN100 for Jurkat cells and EH115 for primary human T cells. Immediately after electroporation, 80 μl of pre-warmed media (without cytokines) was added to each well, and the cells were allowed to rest for 15 min at 37 °C in a cell culture incubator while remaining in electroporation cuvettes. After 15 min, cells were moved to final culture flasks. Jurkat cells were clonally selected by single cell sorting into U-bottomed 96 well plates and by testing each clone using PCR primers flanking the editing site (**Extended Data Fig. 5a**). The clones producing a single PCR band of 1283 bp and 1022 bp were selected as clonal populations for *TCRA* Δ*PAR* and *TCRB* Δ*PAR* respectively.

Genome edited populations of primary T cells with the *TCRA* Δ*PAR* and *TCRB* Δ*PAR* mutations were determined by measuring the density of the PCR bands described above resulting from the edited cell population compared to the PCR band from non edited cells, using ImageJ. For comparison with the *TCRA* Δ*PAR* or *TCRB* Δ*PAR* primary T cell populations, we edited cells from both donors using each gRNA targeting the *TCRA* 3’-UTR and *TCRB* 3’-UTR region separately (single gRNA experiments), a gRNA targeting the coding sequence (CDS) of *TCRA* which knocks out TCR expression with high efficiency (Δ*TCR*)^62^, a scrambled gRNA (SC) which does not target any site in the human genome, and cells mixed with the gRNA/Cas9 RNPs but not nucleofected.

### Western Blot

Western blot analysis was performed using the following antibodies: anti-TCRA (SC-515719), anti-HSP90 (SC-69703) (**Supplementary Table 7**).

### Total mRNA isolation and quantitative RT-PCR analysis

Total RNA was isolated from whole cells for qRT-PCR using Quick RNA miniprep plus kit from Zymo Research following the manufacturer’s instructions. Quantitative RT-PCR analysis was performed using the Power SYBR Green RNA-to-Ct 1-Step kit (Applied Biosystems) according to the manufacturer’s instructions, and the QuantStudio™ 3 Real-Time PCR System (ThermoFisher). Each target mRNA was quantified in three biological replicates, with each biological replicate having three technical replicates.

### Plasmids

Nanoluciferase reporters^63^ were constructed using the 5’-UTR of the human beta globin mRNA (*HBB*) and a PEST destabilization domain. The PEST domain reduces protein half-life^64^ and was used to provide better time resolution of nanoluciferase expression after T cell activation. The *TCRA* 3’-UTR and *TCRB* 3’-UTR sequences were amplified from Jurkat genomic DNA. The nanoluciferease sequence fused to a PEST domain was amplified from pNL1.2[*NlucP*] Vector Sequence (Promega) and was cloned into a modified CD813A vector (System Biosciences) containing the beta-globin (*HBB*) 5’-UTR using the In-Fusion® HD Cloning Kit (Takara). The subsequent mutations in the *TCRA* and *TCRB* 3’-UTRs were generated using these initial constructs. For *TCRA* Δ*PAR* constructs, nucleotides 102-338 in the 3’-UTR of *TCRA* mRNA were deleted. For *TCRB* Δ*PAR* constructs, nucleotides 16-161 in the 3’-UTR of *TCRB* mRNA were deleted. *TCRA/TCRB* Δ*PAR, TCRA/TCRB R*PAR, 3’-LTR* (3’-Long Terminal Repeat) and *bpA* (bovine growth hormone polyadenylation signal) sequences were purchased as gblocks from IDT and were cloned into this plasmid backbone. The *WPRE* 3’-UTR sequence was amplified from the CD813A-1 (System Biosciences) vector.

For nanoluciferase reporters designed to be membrane-tethered, we fused the N-terminal se quence of CD3-zeta spanning the transmembrane helix (amino acids 1-60) to the nanoluciferase sequence above. To prevent interaction of the CD3-zeta-nanoluciferase fusion protein with the TCR, we made mutations in the CD3-zeta derived transmembrane helix that would disrupt interactions with the TCR, based on the cryo-EM structure of the complex^65^ (PDB entry 6JXR) and consistent with earlier biochemical results^66^. The two mutations, L35F and D36V, are predicted to introduce a steric clash and disrupt an intra-membrane salt bridge, respectively, with other subunits in the TCR holo-complex. These CD3-zeta-nanoluciferase chimeras were cloned into the modified CD813A plasmids described above.

### Generation of stable Jurkat cell lines expressing nanoluciferase reporters

For lentiviral production, HEK293T cells were plated at a density of 80% in T-75 flasks the night before transfection. The cells were then transfected with plasmids: expressing the nanoluciferase, PsPAX2 and pCMV-VSV-G using the Lipofectamine 3000 reagent (ThermoFisher) following the manufacturer’s instructions. Forty-eight hours after transfection, the viral supernatant was collected, filtered and concentrated using PEG-it Virus Precipitation Solution (System Biosciences) following the manufacturer’s instructions. The virus pellets were then resuspended in RPMI-1640 media and stored in -80 °C.

The Jurkat cell transductions were done with multiple viral titers using TransDux™ MAX Lentivirus Transduction Reagent (System Biosciences) following the manufacturer’s instructions. To test the viral transduction efficiency of the cells, forty-eight hours after viral transduction the percent of cells expressing GFP was measured by FACS analysis and cells expressing less than 15% GFP were treated with 1 µg ml^-1^ puromycin to the media. The cells were maintained in media with 1 µg ml^-1^ puromycin, but transferred into media without puromycin the day before the cells were used for assays.

### Luciferase reporter assay

The stable cell lines expressing the Nanoluciferase reporters were stimulated with anti-CD3 and anti-CD28 antibodies and the nanoluciferase activity was assayed after 1 hr, 3 hr, 5 hr, 8 hr, 10 hr and 24 hrs after stimulation using Nano-Glo® Luciferase Assay System (Promega). For each time point 200,000 cells were tested in triplicate for each cell line.

### RNA immunoprecipitation and qPCR

The EIF3B-RNA immunoprecipitations were carried out following the exact same conditions used for the PAR-CLIP analysis with the following changes. For each immunoprecipitation, cell lysates were prepared in NP40 lysis buffer (defined in the PAR-CLIP section) with 4 million cells. The lysates were then incubated with 50 μl Protein G

Dynabeads conjugated with 20 μl of anti-EIF3B antibody (Bethyl A301-761A) for two hours at 4 °C. After incubation, the flow through was removed and the beads were washed three times with 1 ml NP40 lysis buffer for each wash. The beads were then resuspended in 400 μl TRIzol reagent (ThermoFisher) and vortexed for 1 minute. The RNA was extracted following the manufacturer’s instructions and qPCR was performed as described above using primers listed in **Supplementary Table 7**.

### Flow cytometry and cell sorting

Flow cytometric analysis was performed on an Attune NxT Acoustic Focusing Cytometer (ThermoFisher). Surface staining for flow cytometry and cell sorting was performed by pelleting cells and resuspending in 50 μl of FACS buffer (2% FBS in PBS) with antibodies at the indicated concentrations (**Supplementary Table 7**) for 20 min at 4 °C in the dark. Cells were washed twice in FACS buffer before resuspension.

### ELISA

The cell suspensions were collected after each time point of activation with anti-CD3/anti-CD28 antibodies for WT, *TCRA* Δ*PAR* or *TCRB* Δ*PAR* cells. For each timepoint the same number of cells were used from each strain to be able to compare across strains and time points. The amount of secreted IL2 in the cell suspensions after activation anti-CD3/anti-CD28 antibodies for WT, *TCRA* Δ*PAR* or *TCRB* Δ*PAR* cells were measured by ELISA MAX™ Deluxe Set Human IL-2 (BioLegend) according to the manufacturer’s instructions.

### RNA-FISH and immunofluorescence

Jurkat cells were washed with PBS, fixed with 3.7% (vol./vol.) formaldehyde for 10 min at room temperature and washed three times with PBS. PBS was discarded and 1 ml 70% ethanol was added. The cells were incubated at 4 °C for 16 hours. The 70% ethanol was aspirated and the cells were washed once with 0.5 ml Stellaris RNA wash buffer A (Biosearch technologies). The cells were then incubated with 100 μl Stellaris hybridization buffer (Biosearch Technologies) containing Stellaris RNA FISH probes (Biosearch Technologies) at a final concentration of 125 nM (**Supplementary Table 7)** and with the relevant antibody (**Supplementary Table 7)** for 16 hours at 28 °C. The cells were then washed twice with 0.5 ml Stellaris RNA wash buffer A containing secondary antibody conjugated with a fluorophore for 30 minutes at 37 °C in the dark. The second Stellaris RNA wash buffer A contained DAPI in addition to the secondary antibody. Finally the cells were washed once with 0.5 mL Stellaris RNA wash buffer B and mounted with mounting solution (Invitrogen). All high resolution images were taken using confocal ZEISS LSM 880 in Airyscan super-resolution mode, equipped withA Plan-Apochromat 63x/1.4 Oil objective (Zeiss). To measure colocalization of *TCRA* and *TCRB* mRNAs with each other or with P bodies (using DCP1 antibody) or stress granules (G3BP1 antibody) the cells were mounted along with 0.1 µm TetraSpeck™ microspheres (ThermoFisher) adhered to the slide according to manufacturer’s instructions, to be able to account for the chromatic shift during image acquisition. For the counting of cells containing TCR clusters, a Revolve Epi-Fluorescence microscope (Echo), equipped with an A Plan-Apochromat 40x objective (Olympus) was used.

### Colocalization analysis

To measure colocalization of *TCRA* and *TCRB* mRNAs with each other or with P bodies (using DCP1 antibody) or stress granules (G3BP1 antibody), immunofluorescently labelled cells (see above) were mounted along with 0.1 µm TetraSpeck™ microspheres (ThermoFisher) adhered to the slide according to manufacturer’s instructions. The microspheres allowed for the correction of lateral and axial chromatic aberrations during image acquisition.

Z-stacks were acquired with 35 nm x 35 nm x 190 nm voxels on a ZEISS LSM 880 in Airyscan super-resolution mode, equipped with a Plan-Apochromat 63x/1.4 Oil objective (Zeiss). The images were then deconvolved to a lateral resolution of ≈150 nm and an axial resolution of ≈500 nm (as confirmed by observing the discrete Fourier transform of the z-stacks). After imaging a single cell, beads that were on the slide axial to the cell were imaged to measure the corresponding lateral and axial chromatic aberrations.

To quantify the relative colocalization, we developed an automated processing and analysis pipeline in ImageJ 1.52p available on github: https://github.com/Llamero/TCR_colocalization_analysis-macro. Specifically, the chromatic aberrations in the z-stacks were compensated for by registering the channels of the bead z-stack to one another, and then applying the same registration vectors to the corresponding channels in the cell z-stack^67^. Each channel of a z-stack was then thresholded to remove background in the image, and then the colocalization between each pair of images was measured using the Pearson’s correlation coefficient. Samples in which any pair of channels in the bead z-stack had a correlation of less than 0.45 were removed from final analysis, as this suggested that the images had insufficient dynamic range in at least one of the channels for an accurate deconvolution.

## Data and Code Availability

Sequencing data supporting the findings in this study have been deposited in the Gene Expression Omnibus (GEO) Database (https://www.ncbi.nlm.nih.gov/geo/), with accession codes. Code used to analyze the microscopy images is available on github at https://github.com/Llamero/TCR_colocalization_analysis-macro. Raw data for **Figure 2a, Figure 3a, Figure 4a**, and **Extended Data Fig. 1a-c** are provided in Supplementary Figures 1-3.

### Acknowledgements

We thank M. Hafner for advice on PAR-CLIP methodology and data analysis, A. Weiss for experimental suggestions and advice, H. Nolla and A. Valeros at the UC Berkeley flow cytometry facility for helping out with the FACS analysis and single cell sorting, F. Ives and H. L. Aaron at the UC Berkeley Imaging center for help with microscopy, J. Lui and J. Bohlen for advice on planning experiments and for suggestions on the manuscript, N. Aleksashin and A.M. González-Sánchez for advice on the manuscript, and M. Mignardi for advice on FISH experiments. This work was supported by NIH grant R01-GM065050 (J.H.D.C.) and a grant from the Tang Prize for Biopharmaceutical Science (J.H.D.C.). N.T.I. is a Damon Runyon-Rachleff Innovator (DRR#37-15) and NIH New Innovator (DP2 CA195768). A.M. holds a Career Award for Medical Scientists from the Burroughs Wellcome Fund and The Cancer Research Institute (CRI) Lloyd J. Old STAR grant and has received funding from the Chan Zuckerberg Biohub, Innovative Genomics Institute (IGI) and the Parker Institute for Cancer Immunotherapy (PICI). B.E.S. is supported by NIH grant P30EY003176. Imaging experiments were conducted at the Cancer Research Laboratory Molecular Imaging Center, supported by Helen Wills Neuroscience Institute. Sequencing was carried out at the UCSF Center for Advanced Technology and at the Vincent J. Coates Genomics Sequencing Laboratory at UC Berkeley, supported by NIH grant S10 OD018174.

## Author Contributions

Project and experiments were conceived and designed by D.D.S. and J.H.D.C. D.D.S. carried out all experiments, with assistance from B.E.S. and G.H.C. for image acquisition and analysis, from T.R., R.A.A., and A.M. for primary T cell genome editing and cell culture protocols, and from L.F., M.K., N.T.I. and J.H.D.C. for analysis of the deep sequencing-based experiments. The manuscript was written by D.D.S. and J.H.D.C., with editing by all authors.

## Competing Interests

The authors declare competing financial interests: T.L.R. and A.M. are co-founders of Arsenal Therapeutics. A.M. is a co-founder of Spotlight Therapeutics. A.M. served on the scientific advisory board of PACT Pharma, was an advisor to Juno Therapeutics and Trizell. The Marson Laboratory has received research support from Juno Therapeutics, Epinomics, Sanofi, GlaxoSmithKline,Gilead and Anthem. A provisional patent application has been filed on some of the work presented herein.

## Materials and Correspondence

Correspondence and requests for materials should be addressed to J.H.D.C..

## Extended Data Figures

**Extended Data Figure 1.**
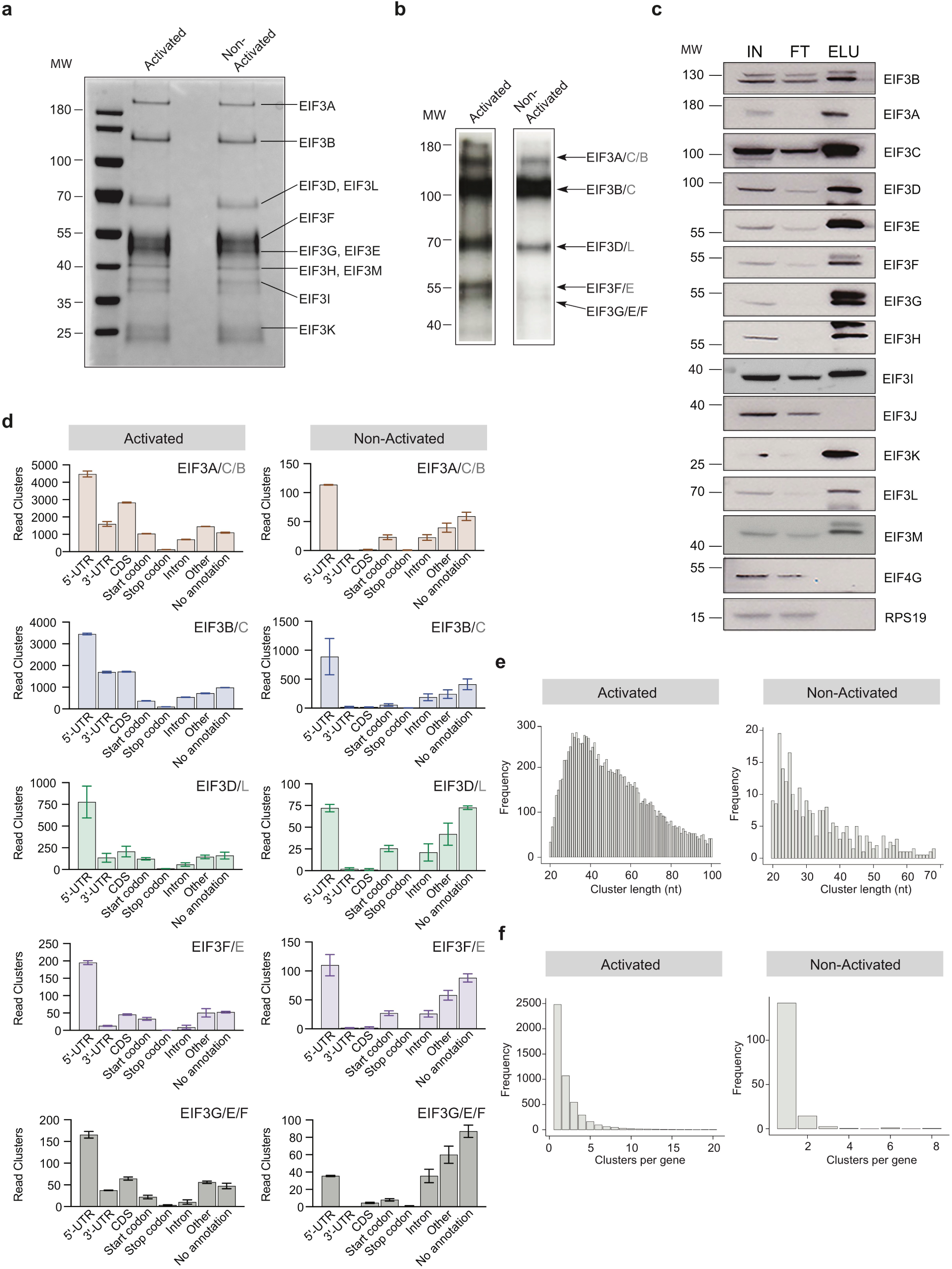
Composition of eIF3 and eIF3-crosslinked mRNAs in the Jurkat PAR-CLIP experiments. **a**, Composition of eIF3 in I+PMA activated and non-activated Jurkat cells after anti-EIF3B immunoprecipitation (IP), identified by mass spectrometry. Shown is an SDS polyacrylamide gel stained with Coomassie Brilliant Blue. **b**, Phosphorimage of SDS polyacrylamide gel resolving 5′ ^32^P-labeled RNAs crosslinked to eIF3 subunits in activated and non-activated Jurkat cells, from one of two biological replicates. **c**, Composition of eIF3 in activated Jurkat cells determined by western blot after anti-EIF3B IP. IN: input; FT: flow-through from anti-EIF3B IP beads; ELU: elution of eIF3 from anti-EIF3B IP beads. Anti-EIF4G1 and anti-RPS19 western blots confirm the stringency of bead wash steps^1^. **d**, Categories of RNA crosslinked to eIF3 by PAR-CLIP in Jurkat cells, defined by clusters and divided into RNA categories, in I+PMA activated Jurkat cells (left) and non-activated Jurkat cells (right). In all panels, RNA categories include: 5’-UTR, 5’-UTR of mRNA; 3’-UTR, 3’-UTR of mRNA; CDS, protein coding region of mRNA; Start codon, beginning of mRNA CDS; Stop codon, stop codon region of mRNA; Intron, regions of pre-mRNA; Other, other classes of RNA; No annotation, reads that map to the human genome but that have no annotation assigned. Clusters in different mRNA categories may occur in the same mRNA species. **e**, The length distribution of PAR-CLIP clusters mapped to RNAs crosslinked to eIF3 in the EIF3A/C/B samples, from activated (left) and non-activated (right) Jurkat cells. **f**, The frequency distribution of the number of eIF3 PAR-CLIP clusters mapped to RNAs in the EIF3A/C/B samples, from activated (left) and non-activated (right) Jurkat cells. In all panels, the distributions are the average of both biological replicates. RNA PAR-CLIP properties observed in the EIF3A/C/B sample are representative of those seen in PAR-CLIP sequence reads of RNAs crosslinked to other eIF3 subunits.

**Extended Data Figure 2.**
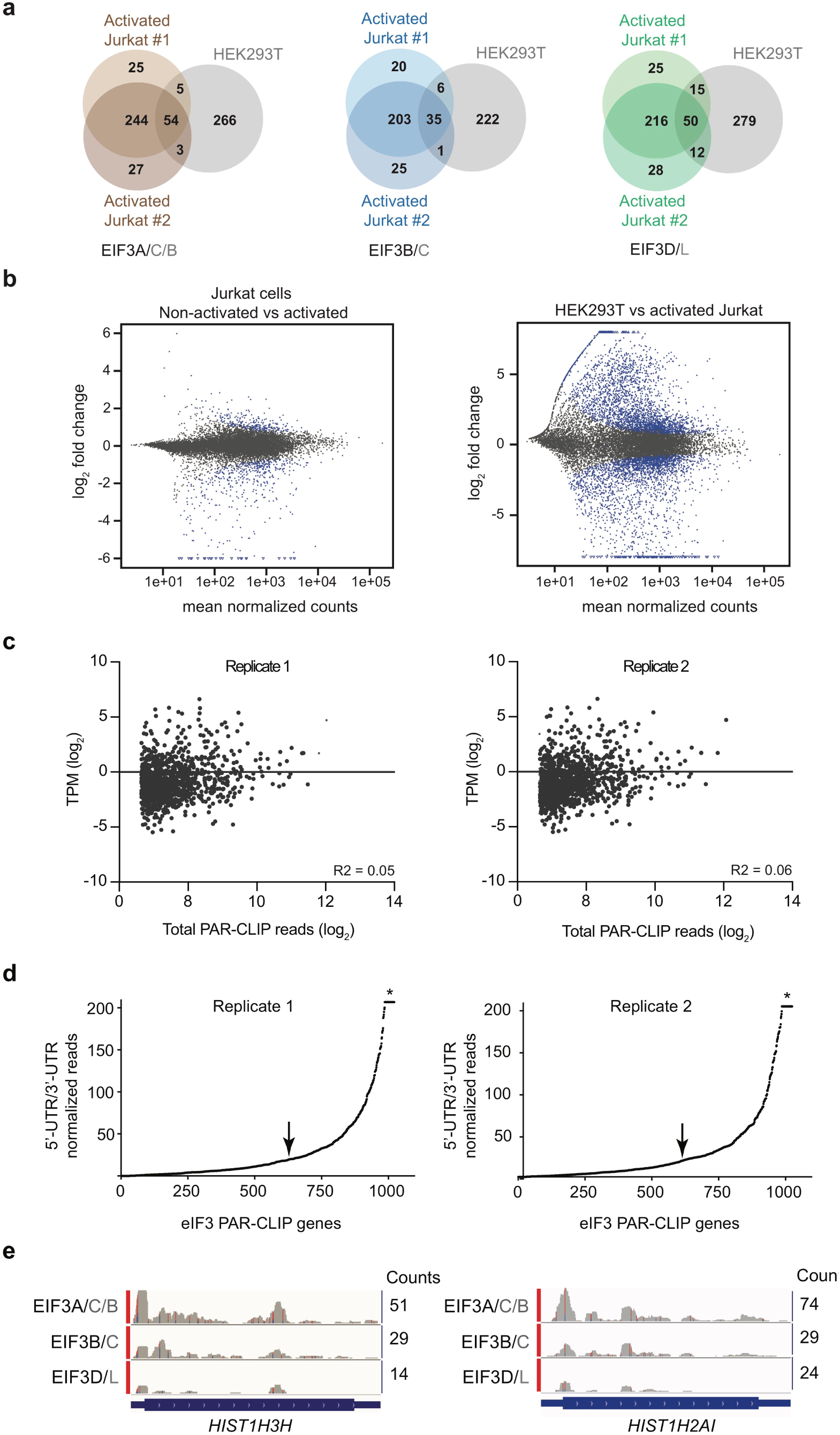
Profiles of eIF3 PAR-CLIP reads in Jurkat cells. **a**, Venn diagram of genes identified by eIF3 PAR-CLIP in the two biological replicates of activated Jurkat cells, compared with the eIF3 PAR-CLIP mRNAs identified in HEK293T cells. The same number of mRNAs from the Jurkat cell PAR-CLIP libraries, ranked by total reads mapped to a given gene, are compared to the number of mRNAs identified in the HEK293T PAR-CLIP experiments. **b**, MA plots of non-activated Jurkat cells compared to Jurkat cells activated with I+PMA for 5 hours (left) and HEK293T cells compared to Jurkat cells activated with I+PMA for 5 hours (right). In the HEK293T plot, M = log2(HEK293T mRNA TPM/activated Jurkat mRNA TPM), and A = mean(TPM) for activated Jurkat mRNAs. TPM, transcripts per million. The FDR was set to 0.01, and the blue dots indicate significant fold changes. **c**, Scatterplot of TPM of mRNAs expressed in activated Jurkat cells (most abundant isoform, see Methods) versus mRNAs identified by PAR-CLIP as crosslinked to eIF3, with ≥ 100 total read counts. n=1,029 and 1,035 for PAR-CLIP hit genes plotted for replicates 1 and 2, respectively. The R-squared goodness of fit to a linear equation is shown. **d**, mRNAs sorted by increasing value of the ratio of normalized 5’-UTR reads to normalized 3’-UTR reads. Values of the ratio of 5’-UTR/3’-UTR normalized reads > 200 have been truncated to a value of 200 in the plot (asterisk). The arrow indicates the threshold used to create cumulative plots shown in **Fig. 1b. e**, Examples of eIF3 crosslinking to histone mRNAs *HIST1H2AI* and *HIST1H3H* in activated Jurkat cells are shown. Transcription start sites were determined from the FANTOM5 Database^68^.

**Extended Data Figure 3.**
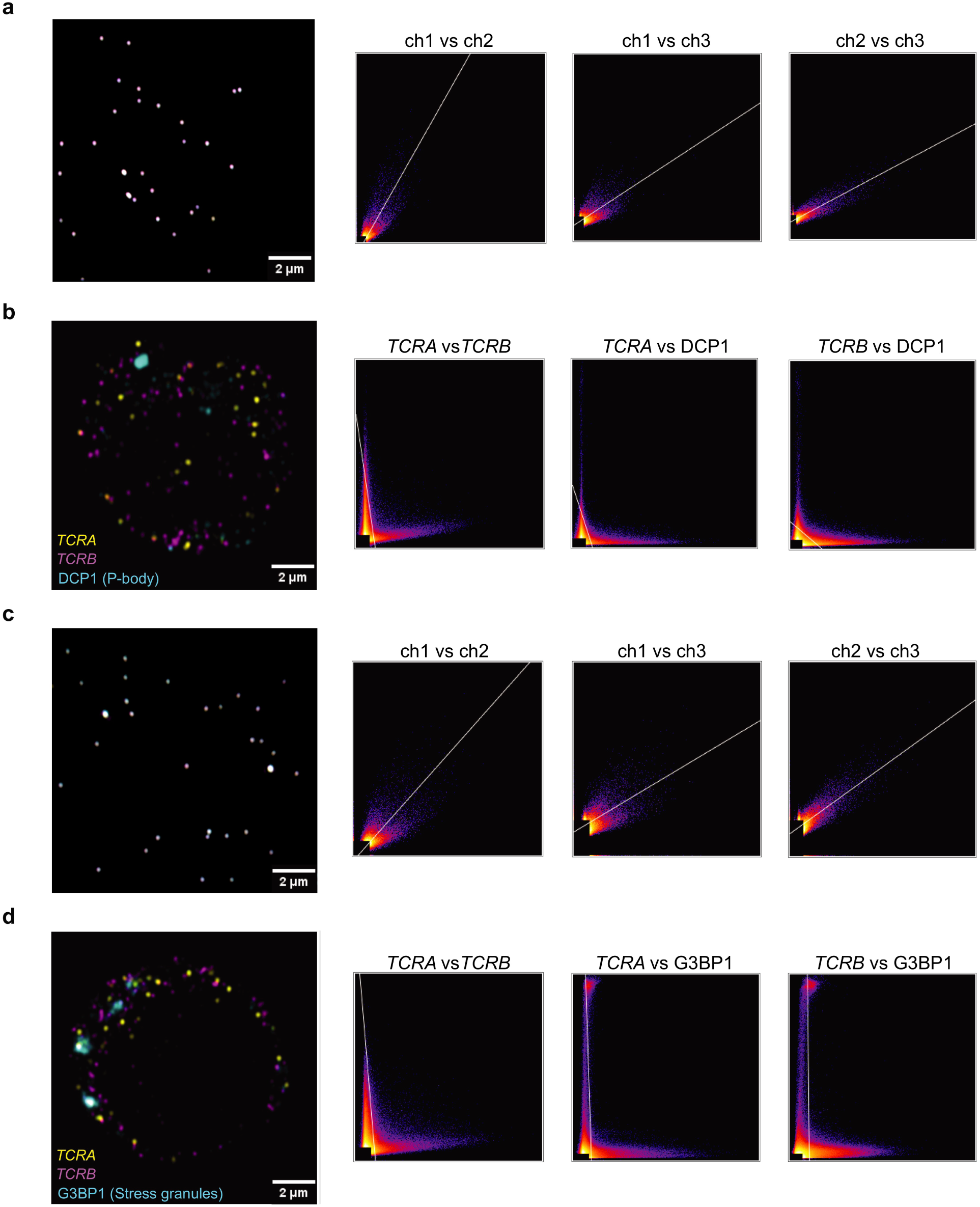
*TCRA* and *TCRB* mRNA cellular localization patterns. **a**, Single optical section of TetraSpeck microspheres (0.1 µm) adjacent to a Jurkat cell probed for P bodies (far left). Scatter plots of pixel intensities in different emission channels are shown: channel 1, 670 nm; channel 2, 570 nm; channel 3, 488 nm. Plots left to right are: channel 1 vs. channel 2, channel 1 vs. channel 3, and channel 2 vs. channel 3. **b**, Single optical section of a Jurkat cell probed for *TCRA* mRNA (channel 1), *TCRB* mRNA (channel 2), and DCP1 to mark P bodies (channel 3). Scatter plots to the right are shown as in **a. c**, Single optical section and scatter plots for TetraSpeck microspheres as in **a**, but adjacent to a Jurkat cell probed for stress granules. **d**, Single plane image of a Jurkat cell probed for *TCRA* mRNA (channel 1), *TCRB* mRNA (channel 2), and G3BP1 to mark stress granules (channel 3). Scatter plots to the right are shown as in **a**. Images of each cell are representative of those used in the analysis in **Figure 1d**.

**Extended Data Figure 4.**
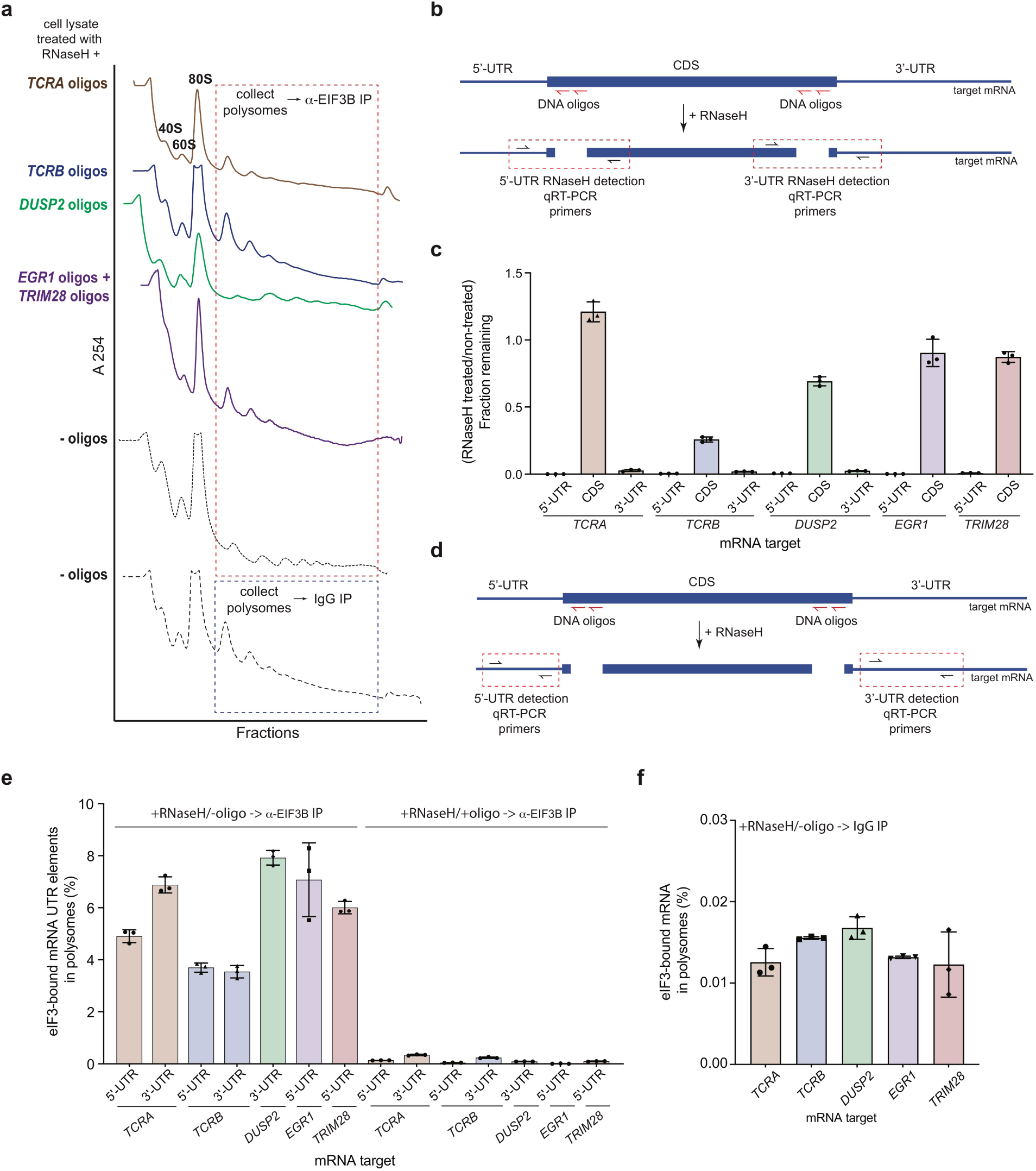
Interactions of eIF3 with mRNAs during translation elongation in activated Jurkat cells. **a**, Sucrose gradient fractionation of polysomes from crosslinked Jurkat cells. Cells lysates treated with RNaseH, as indicated, were fractionated on 10%-50% sucrose gradients. Shown is the absorbance at 254 nm, for one of two biological replicates. **b**, Strategy for detecting mRNA cleavage by RNaseH digestion. RT-qPCR primers were designed to span the mRNA digestion sites. **c**, Fraction of intact mRNA segments remaining after RNaseH treatment of DSP-crosslinked cell lysates, in the presence or absence of mRNA-specific DNA oligos for the indicated mRNAs. The mRNAs were detected using RT-qPCR oligos as illustrated in panel **b. d**, Strategy for detecting 5’-UTR or 3’-UTR mRNA fragments released by RNaseH digestion. RT-qPCR primers were designed within the 5’-UTR or 3’-UTR regions. **e**, Amount of eIF3-bound 5’-UTR and 3’-UTR regions of the mRNA co-immunoprecipitated by anti-EIF3B antibody, from polysome fractions of lysate treated with RNaseH, either in the absence (left) or presence (right) of mRNA-targeting DNA oligos. Percentage is relative to mRNA present in the polysome fraction prior to the IP. Primers to the mRNA 5’-UTR and 3’-UTR regions, as indicated in panel **d**, were used for quantification. **f**, Amount of eIF3-bound mRNA co-immunoprecipitated with IgG beads, from polysome fractions of lysate treated with RNaseH, but in the absence of mRNA-targeting DNA oligos. Percentage is relative to mRNA present in the polysome fraction prior to the IP. All experiments in panels **c, e**, and **f** were carried out in biological duplicate with one technical triplicate shown (*n* = 3, with mean and standard deviations shown).

**Extended Data Figure 5.**
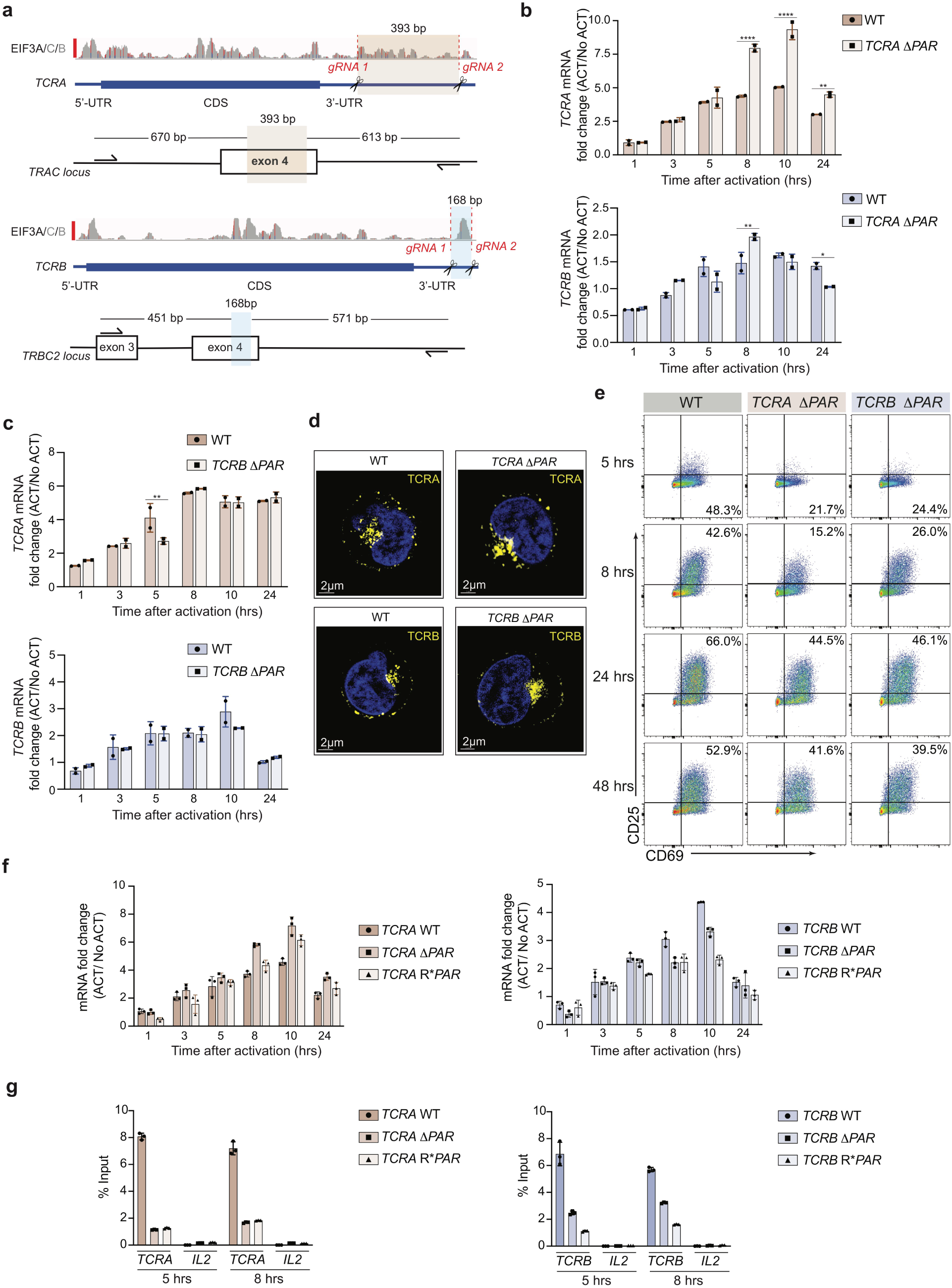
Deletion of eIF3 PAR-CLIP sites in the 3’-UTRs of *TCRA* and *TCRB* mRNAs. **a**, CRISPR/Cas9 RNP mediated genome editing at the *TCRA* and *TCRB* genomic loci in Jurkat and primary human T cells. Location of gRNA sites targeting the *TCRA* 3’-UTR, to generate *TCRA* Δ*PAR* cells. In *TCRA*, gRNA1 and gRNA2 target hg38 genomic locations chr14:22,551,700 and chr14:22,552,073, respectively. **b**, *TCRA* and *TCRB* mRNA levels in WT Jurkat cells, or Jurkat cells with the *TCRA* eIF3 3’-UTR PAR-CLIP site deleted, as determined by qRT-PCR at different time points after anti-CD3/anti-CD28 activation. **c**, *TCRA* and *TCRB* mRNA levels in WT Jurkat cells, or Jurkat cells with the *TCRB* eIF3 3’-UTR PAR-CLIP site deleted, as determined by qRT-PCR at different time points after anti-CD3/anti-CD28 activation. (*n = 2*, with mean ± standard deviations shown). In panels **b** and **c**, **** P < 0.0001, *** P < 0.001, ** P < 0.01 for *TCRA* Δ*PAR* or *TCRB* Δ*PAR* relative to WT, using two-way ANOVA. **d**, TCR clustering in Jurkat cells, after 8 hours of activation with anti-CD3/anti-CD28 antibodies. WT, *TCRA* Δ*PAR, TCRB* Δ*PAR* cells exhibiting TCR clustering, as determined using anti-TCRA/anti-TCRB protein staining and Airyscan confocal microscopy. (*n = 5* for each time point after activation for all cell lines, with representative images shown.) **e**, Density plots showing cell populations expressing CD69+ only, CD69+ CD25+ or CD25+ only from WT, *TCRA* Δ*PAR* or *TCRB* Δ*PAR* Jurkat cells at different time points after stimulation with anti-CD3/anti-CD28 antibodies, (*n = 3*, with representative plots shown). **f**, Changes in nanoluciferase mRNA levels in the experiment shown in **Fig. 2h**, as determined by qRT-PCR. P < 0.0001 for *TCRA* Δ*PAR* relative to WT at 8 hr, and for *TCRA* Δ*PAR* and R\**PAR* relative to WT at 10 hr, using two-way ANOVA. P ≤ 0.0002 for *TCRB* Δ*PAR* and R\**PAR* relative to WT at 8 hr and 10 hr. **g**, Immunoprecipitation of eIF3 using an anti-EIF3B antibody^1^, followed by qRT-PCR to quantify the amount of nanoluciferase reporter mRNA bound to eIF3, for the 5 hour and 8 hour timepoints in the experiment shown in **Fig. 2h**. The percent mRNA bound to anti-EIF3B beads is calculated relative to total mRNA isolated from the cells. (*n* = 3, with mean and standard deviations shown).

**Extended Data Figure 6.**
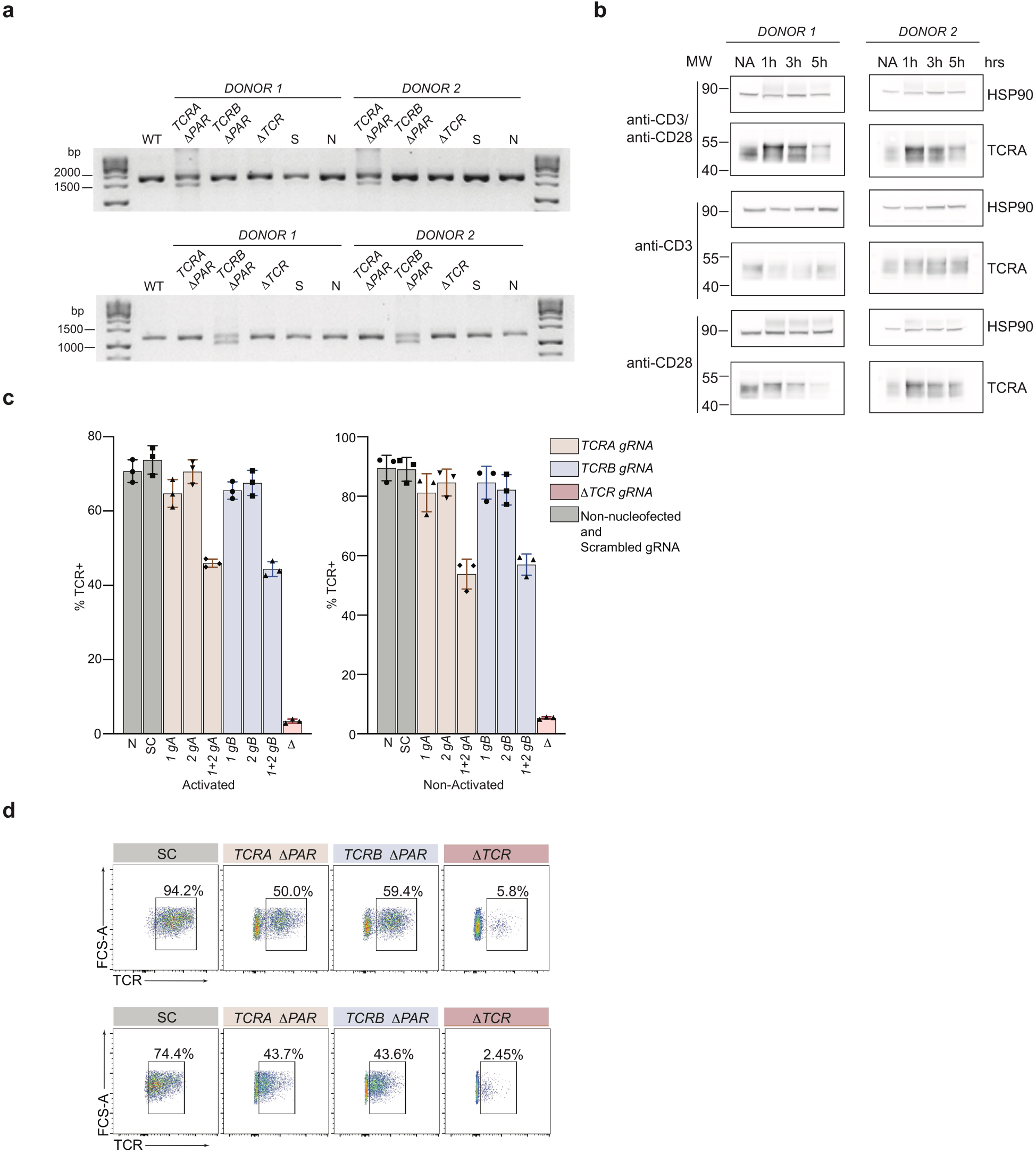
Phenotypic analysis of primary T cells. **a**, Top gel: Analysis of *TCRA* 3’-UTR PAR-CLIP site deletion efficiency. Total genomic DNA extracted from WT, *TCRA* Δ*PAR, TCRB* Δ*PAR* (here serving as a negative control), scrambled gRNA nucleofected (S) and non-nucleofected (N) cells was analyzed by PCR to measure the editing efficiency. *TCRA* Δ*PAR* cells produced a 1283 bp PCR product compared to 1676 bp in S, N or *TCRB* Δ*PAR* cells. Bottom gel: Analysis of *TCRB* 3’-UTR PAR-CLIP site deletion efficiency. Total genomic DNA extracted from WT, *TCRA* Δ*PAR* (here serving as a negative control), *TCRB* Δ*PAR*, scrambled gRNA nucleofected (S) and non-nucleofected (N) cells were analyzed by PCR to measure the editing efficiency. *TCRB* Δ*PAR* cells produced a 1022 bp PCR product compared to 1190 bp in S, N or *TCRA* Δ*PAR* cells. **b**, Western blots of TCRA protein levels in primary human T cells using different activation modes. HSP90 was used as a loading control. **c**, Percentage of cells expressing TCR on the cell surface measured by flow cytometric analysis (*n = 2* donors). TCR expressing cells in the cell populations tested: *N*, Non-nucleofected cells; SC, Scrambled gRNA; *1gA, TCRA* gRNA 1; *2gA, TCRA* gRNA 2; *1+2 gA, TCRA* gRNA 1+2 (i.e. *TCRA* Δ*PAR*); *1gB, TCRB* gRNA 1; *2gB, TCRB* gRNA 2; *1+2 gB, TCRB* gRNA 1+2 (i.e. *TCRB* Δ*PAR*); Δ*TCR*, TCR gRNA targeting the CDS of *TCRA*. (*n* = 2 donors, with standard deviation shown.) Left: I+PMA activated T cells, after 3 hours of activation; Right: non-activated T cells. P values for *TCRA* Δ*PAR, TCRB* Δ*PAR*, Δ*TCR* were < 0.0001 relative to SC in both activated and non-activated cells, using one-way ANOVA. The P values for single guide edited non-activated cells relative to SC were not significant, while in activated cells P = 0.02 – 0.05 relative to SC. **d**, Representative density plots showing the percentage of TCR on the cell surface in non-activated (top) and activated (bottom) T cells for the two donors. The cell lines shown are: SC, *TCRA* Δ*PAR, TCRB* Δ*PAR*, Δ*TCR* (negative control).

**Extended Data Figure 7.**
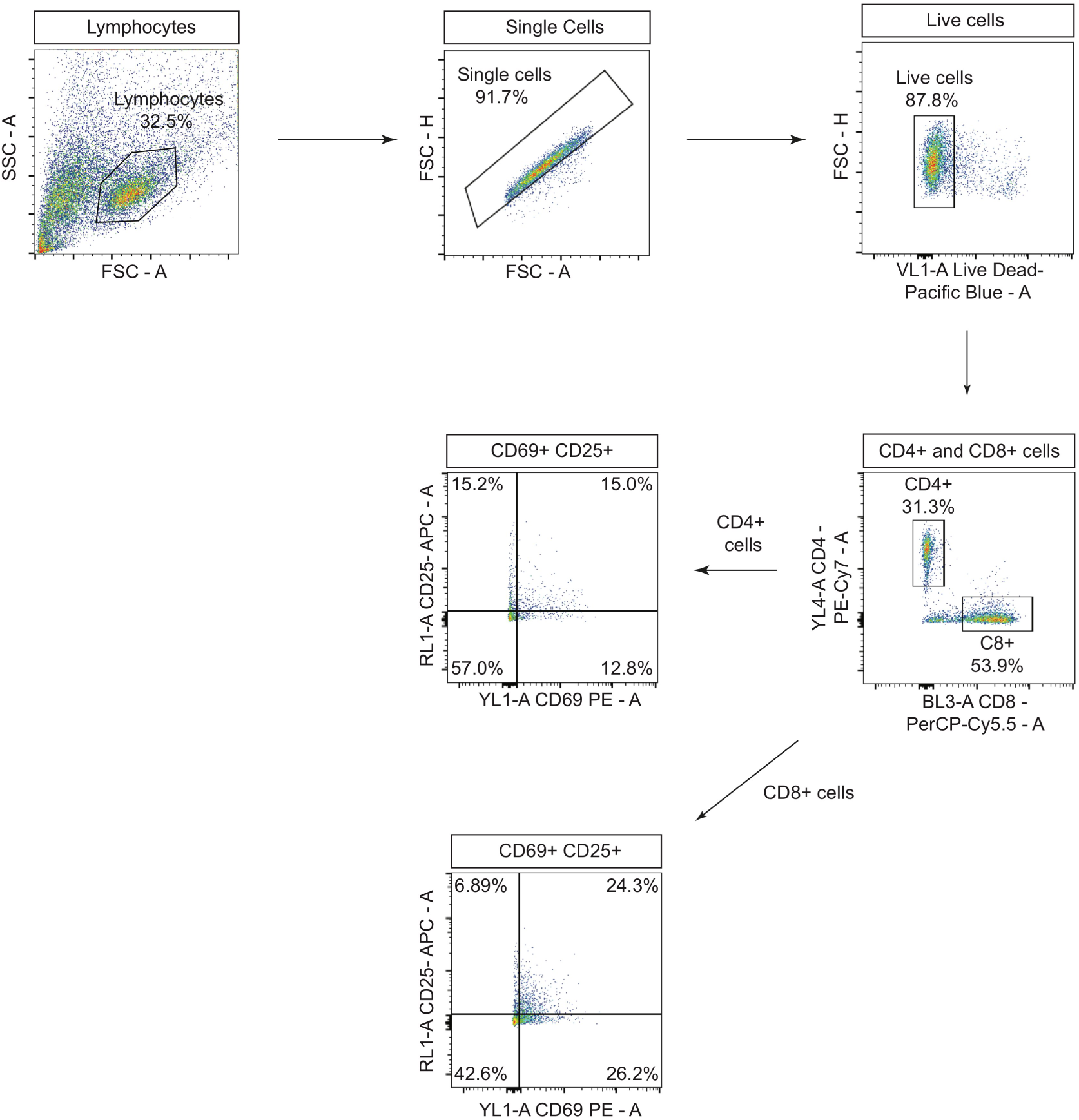
Gating strategy for flow cytometric analysis of CD69 and CD25 levels. Primary human T cells were gated to isolate lymphocytes, followed by isolation of single cells, then live cells. Next, cells were gated to separate CD4+ and CD8+ cells expressing T cell activation markers CD69 and CD25 after activation with anti-CD3/anti-CD28 antibodies. Shown is the workflow of the FACS gating, with an example of T cells activated with anti-CD3/anti-CD28 antibodies for 8 hours.

**Extended Data Figure 8.**
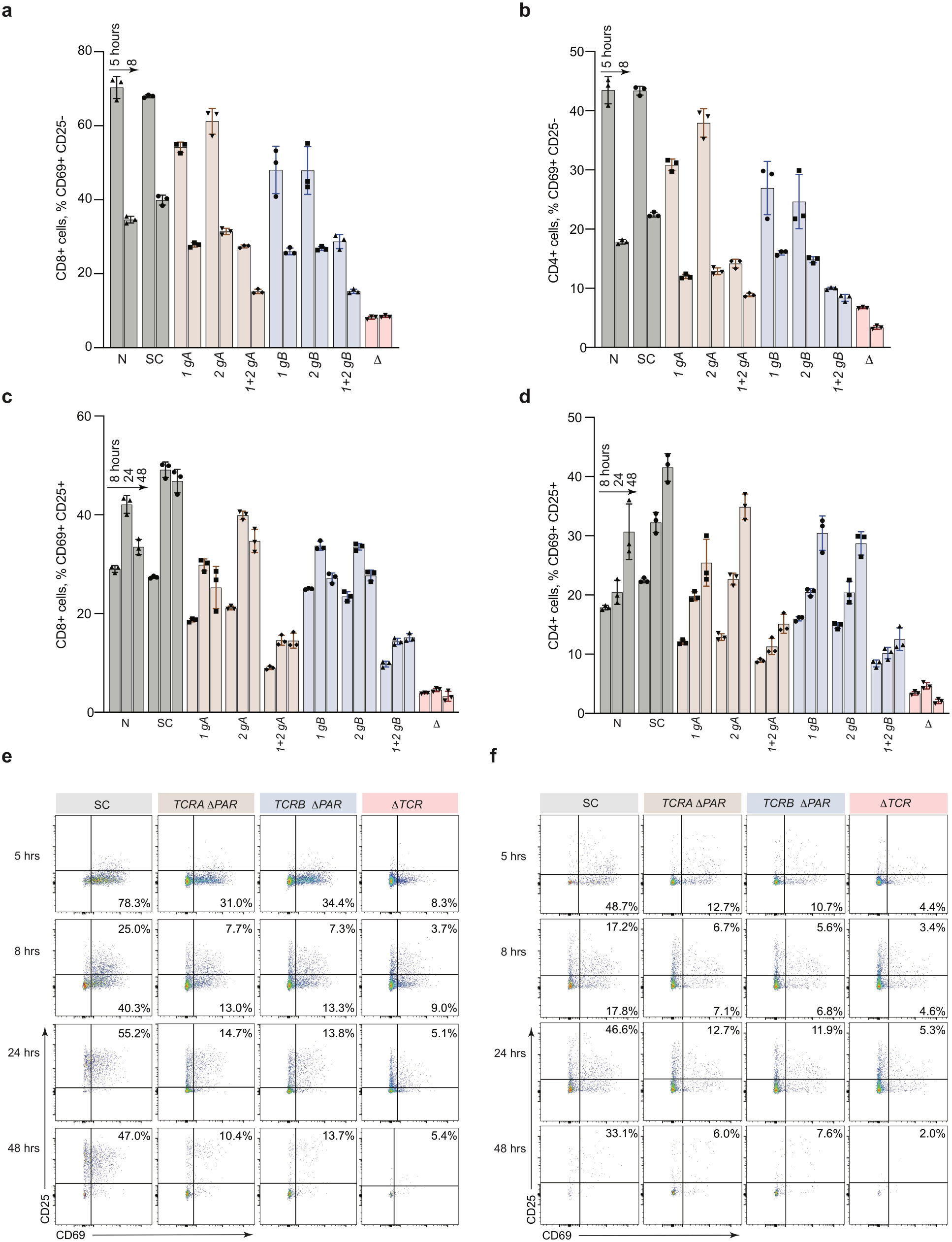
Phenotypic analysis of Primary human T cells with eIF3 3’-UTR PAR-CLIP sites deleted. **a-d**, Flow cytometric analysis of CD8+ and CD4+ T cells showing quantification of T cell activation markers CD69 (early activation marker) and CD25 (mid-activation marker) after activation with anti-CD3/anti-CD28 antibodies at different time points, for all cell populations tested: N, Non-nucleofected cells; SC, Scrambled gRNA; *1gA, TCRA* gRNA 1; *2gA, TCRA* gRNA 2; *1+2 gA, TCRA* gRNA 1+2 (i.e. *TCRA* Δ*PAR*); *1gB, TCRB* gRNA 1; *2gB, TCRB* gRNA 2; *1+2 gB, TCRB* gRNA 1+2 (i.e. *TCRB* Δ*PAR*); *ΔTCR* gRNA targeting the CDS of *TCRA*. (*n = 2* donors, with mean and standard deviation shown). In CD8+ T cells P values for *TCRA* Δ*PAR, TCRB* Δ*PAR*, Δ*TCR* were < 0.0001 relative to SC, while single gRNA edited cell lines had P < 0.0001 to P < 0.01, relative to SC as calculated by two-way ANOVA. In CD4+ T cells P values for *TCRA* Δ*PAR, TCRB* Δ*PAR*, Δ*TCR* were < 0.0001 relative to SC, while single gRNA edited cell lines had P < 0.0001 to P < 0.01, relative to SC as calculated by two-way ANOVA. Representative density plots showing the percentage of **e**, CD8+ T cells and **f**, CD4+ T cells expressing CD69, CD69 and CD25 or CD25 only, for one of two donors. The cell lines shown are SC, *TCRA* Δ*PAR, TCRB* Δ*PAR* and Δ*TCR* (negative control).

**Extended Data Figure 9.**
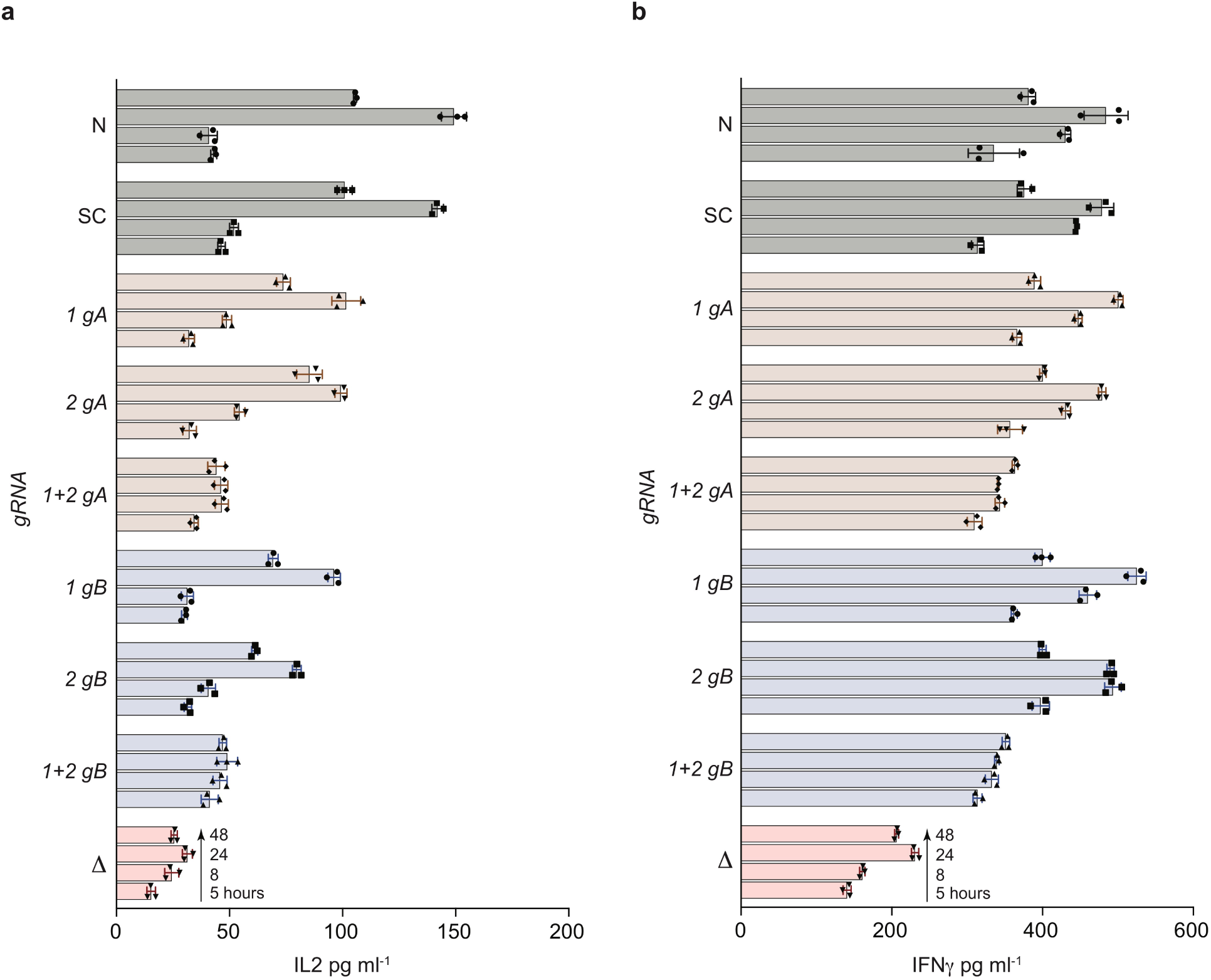
Cytokine secretion profile of primary human T cells with eIF3 3’-UTR PAR-CLIP sites deleted. Quantification of **a**, secreted IL2 and **b**, secreted IFNγ, for all cell populations tested, as determined by ELISA. Cell populations tested were: N, Non-nucleofected cells; SC, Scrambled gRNA; *1gA, TCRA* gRNA 1; *2gA, TCRA* gRNA 2; *1+2 gA, TCRA* gRNA 1+2 (i.e. *TCRA* Δ*PAR*); *1gB, TCRB* gRNA 1; *2gB, TCRB* gRNA 2; *1+2 gB, TCRB* gRNA 1+2 (i.e. *TCRB* Δ*PAR*); *ΔTCR* gRNA targeting the CDS of *TCRA*. (*n = 2* donors, with mean and standard deviation shown.) P values for *TCRA* Δ*PAR, TCRB* Δ*PAR*, Δ*TCR* were < 0.0001 relative to SC, while single gRNA edited cell lines had P < 0.0001 to P < 0.01, relative to SC as calculated by two-way ANOVA.

## Supplementary Information

Supplementary Figures 1-3: Raw images of gels used for **Fig. 2a, Fig. 3a, Fig. 4a** and **Extended Data Fig. 1a-c**.

Supplementary Table 1: Subunits in eIF3 crosslinked to RNA in activated Jurkat cells. Lists the eIF3 subunits, percent sequence coverage and number of identified peptides.

Supplementary Table 2: PARpipe statistics for eIF3 PAR-CLIP samples. Samples are indexed in the first tab, including both biological replicates for activated and non-activated Jurkat cells. Statistics are given for each library at the read, cluster and group level.

Supplementary Table 3: PAR-CLIP mapping to individual genes for each eIF3 PAR-CLIP sample. Samples are indexed in the first tab, including both biological replicates for activated and non-activated Jurkat cells. First lists the gene name and number of clusters identified. The statistics also include: Sum, sum of that statistic over all sites for that gene; Med, median of that statistic for all sites for that gene; ReadCount, total reads mapping to the gene; T2Cfraction, number of reads with T-to-C conversions / number of reads; ConversionSpecificity, log(number of reads with T-to-C conversions / number of reads with other conversions); UniqueReads, reads collapsed to single copies. Also included: 5’utr/Intron/Exon/3’utr/Start_codon/Stop_codon, number of sites mapping to that annotation category; Junction, number of sites mapping to a junction between categories (coding-intron, coding-3’utr, etc.); GeneType, as described in the gene_type category for this gene in the.gtf file used.

Supplementary Table 4. Transcriptome analysis of HEK293T cells and non-activated or activated Jurkat cells. Each tab lists transcript name and version, gene name, type of transcript, length of transcript, and mean transcripts per million, calculated from two biological replicates.

Supplementary Table 5. Differential expression analysis of HEK293T or non-activated vs. activated Jurkat cells. Each tab lists the gene name, mean of normalized counts, log2-fold change, log2-fold change standard error, Wald statistic, Wald test P-value, and Benjamini-Hochberg adjusted P-value.

Supplementary Table 6. Pathway enrichment analysis. Lists for both biological replicates of the EIF3A/C/B PAR-CLIP libraries are included (genes with >= 100 reads), along with associated transcript names, lengths in nts of the 5’-UTR, coding region, and 3’-UTR, and reads normalized to the lengths of the transcript regions. Tabs also include the top tissue-specific pathway enrichment categories determined using the STRING Database. These list: the Gene Ontology (GO) number, GO description, observed gene count, background gene count, false discovery rate, and matching proteins in the network by Ensembl protein ID, and by gene name.

Supplementary Table 7. Reagent information for experiments. Lists include antibodies used, PCR primers, qPCR primers, gRNA targeting sequences, and FISH probes, and DNA oligos for RNaseH experiments.

## Notes

### Summary of Updates

1.We reorganized the paper to present the mechanistic results more clearly. 2.We now show that eIF3 directly contacts the *TCRA* and *TCRB* mRNA coding sequences (CDS) in polysomes (Fig. 1e-h, Extended Data Fig. 4). 3.Using membrane-tethered nanoluciferase reporters, we show CD28 signaling requires cotranslational membrane insertion of the protein for the *TCRA* 3'-UTR to elicit the burst in translation upon T cell activation (Fig. 3). 4.We updated experiments showing clinically-used 3'-UTRs for CAR-T expression do not elicit a burst in translation upon T cell activation (Fig. 4g).

